# Graph pangenome reveals functional, evolutionary, and phenotypic significance of human nonreference sequences

**DOI:** 10.1101/2022.09.05.506692

**Authors:** Zhikun Wu, Tong Li, Zehang Jiang, Jingjing Zheng, Yun Liu, Yizhi Liu, Zhi Xie

## Abstract

Thousands of DNA sequences in global populations are not present in the human reference genome, named nonreference sequence (NRS). Long-read sequencing (LRS) technologies enable better discovery of NRS with large length, particularly in repetitive regions. Here, we *de novo* assembled 539 genomes in five genetically divergent human populations sequenced by LRS technology and identified 5.1 million NRSs. These NRSs were merged into 45,284 nonredundant NRSs, of which 66.2% were novel. 78.5% of NRSs were repeat sequences, such as VNTR and STR. 38.7% of NRSs were common in the five populations, 35.6% were population specific, while 21.3% were ancestral and present in nonhuman primates. 144 NRS hotspots spanned 141 Mb of the human genome and many NRSs contained known functional domains or intersected with coding genes. Based on graph-based pangenome, we detected 565 transcript expression quantitative trait loci on NRSs, of which 467 were novel. We also detected 39 NRS candidates for adaptive selection within the human population related to the language system and diabetes. GWAS revealed 14 NRSs significantly associated with eight phenotypes, such as anaemia. Furthermore, we identified 154 NRSs in strong linkage disequilibrium with 258 phenotype-associated SNPs in the GWAS catalogue. Our work expands the landscape of human NRS and provides novel insights into functions of NRS to facilitate evolutionary and biomedical research.

## Introduction

The human reference genome provides a coordinate for human sequence alignment and has greatly advanced human genetic research, such as genetic variant discovery, gene-disease association, and population genetics^1^. However, the current reference genome, GRCh38, still has many gaps^2^. Another known limitation of the reference genome is the lack of genetic representation. The human reference was derived from the genomes of just over 20 individuals and the inferred ancestral sequences from African, European and East Asian were 57%, 37% and 6%, respectively^3^. Therefore, the reference genome does not capture the diversity of the world population, particularly Asian, which counts 59.8% of the total world population^4^.

Nonreference sequences (NRSs) are defined as sequences missing from the reference genome but present in a subset of a population^3^. In the past several years, large-scale human sequencing projects have started touching on discovering NRS^3,5^. For example, previous studies have revealed 0.33, 29.5 and 46 Mb NRSs in Icelandic, Chinese and Swedish populations, respectively^6^. However, these studies were mainly based on *de novo* assemblies of short or linked reads derived from next-generation sequencing (NGS) platforms, which are difficult for the assembly of segmental duplications, low-complexity and GC bias regions, particularly for the local assemblies of unmapped reads against the reference genome^7,8^.

Long-read sequencing (LRS) platforms, such as Pacific Biosciences (PacBio) continuous long read (CLR), PacBio high-fidelity (HiFi) and Oxford Nanopore Technologies (ONT), are known to generate high-continuity *de novo* genome assemblies^9^. The advantages of the assembly of repetitive regions based on LRS make it beneficial for discovering NRS with large length and in structurally complex genomic regions^3^. For example, some recent human genomes assembled using LRS data, such as a Chinese genome (HX1)^10^ and two Swedish genomes^11^, had higher contiguity, with contig N50 lengths ranging from 8.3 to 9.5 Mb, and revealed 12.8 and 12.2 Mb of NRSs per individual, respectively. Particularly, two decades after the first human genome release, the first complete sequence of a human genome, known as T2T-CHM13, presents another great milestone of Human Genome Project, where 238 megabases (Mb) of sequence were added and corrected^2^.

More recently, the Human Pangenome Reference (HPR) Consortium has proposed an ambitious project aiming to create a more sophisticated and complete human reference genome with a graph-based, telomere-to-telomere (T2T) representation of global genomic diversity^12^. The human pangenome project has just released the first draft human pangenome reference, including 47 phased, diploid assemblies^13^. Although the study has not focused on NRS discovery, it identified many novel variants, haplotypes and alleles at structurally complex loci. As more individuals were included, the HPR will contain a more complete representation of global genomic variation and NRS and will serve as the ultimate genetic resource for biomedical research and precision medicine^12^.

Despite great progress has been made, how widespread NRSs exist in human genome and in human populations is still unclear. Furthermore, the functional, evolutionary, and phenotypic significance of NRS remains largely unexplored. To answer these questions, we systematically identified NRS from 539 assembled human genomes in five populations sequenced by LRS and constructed a graph pangenome. We identified 5.1 million NRSs in total, which combined into 45,284 high-confidence NRSs with an accumulated length of 59.7 Mb, where 66.2% of NRSs were novel. We characterized distribution of NRS in the human genome, in the human population as well as nonhuman primates. Furthermore, we functionally annotated NRS and explored their functions in evolution and disease. The graph-based pangenome of NRSs had a clear advantage in the reference representation of diverse populations. The pangenome significantly improved the read mapping rate and the power of detecting expression quantitative trait loci (eQTLs). We detected 565 expression quantitative trait loci on NRSs, of which 467 were novel. We also identified 39 NRSs related to the language system, anaemia, and diabetes as signatures of local adaptation in diverse populations. GWAS of NRS identified 14 NRSs significantly associated with eight phenotypes and 154 NRSs in strong linkage disequilibrium (LD, r^2^ > 0.8) with 258 phenotype-associated SNPs of the GWAS catalogue, of which 25 (16.2%) NRSs were validated to be significantly associated with the transcript levels, suggesting that these NRSs served as potential candidate causal genes for diverse phenotypes. Our work provided a framework to construct a graph-based pangenome of NRS from LRS datasets and link them to phenotypes. Our work also provided important genomic resources and novel insights into functions of NRS to facilitate evolutionary and biomedical research.

## Results

### NRS discovery

To identify reliable NRSs from *de novo* assemblies of LRS data, we developed a pipeline that included two major steps: (1) the *de novo* assembly of genomes and (2) the extraction of high-confidence NRSs (**Fig. 1a, 1b** and **Methods**). Combined with the construction of graph genome step described in the later section, we provided a framework to build a graph pangenome of NRSs, named GraphNRS.

**Figure 1.**
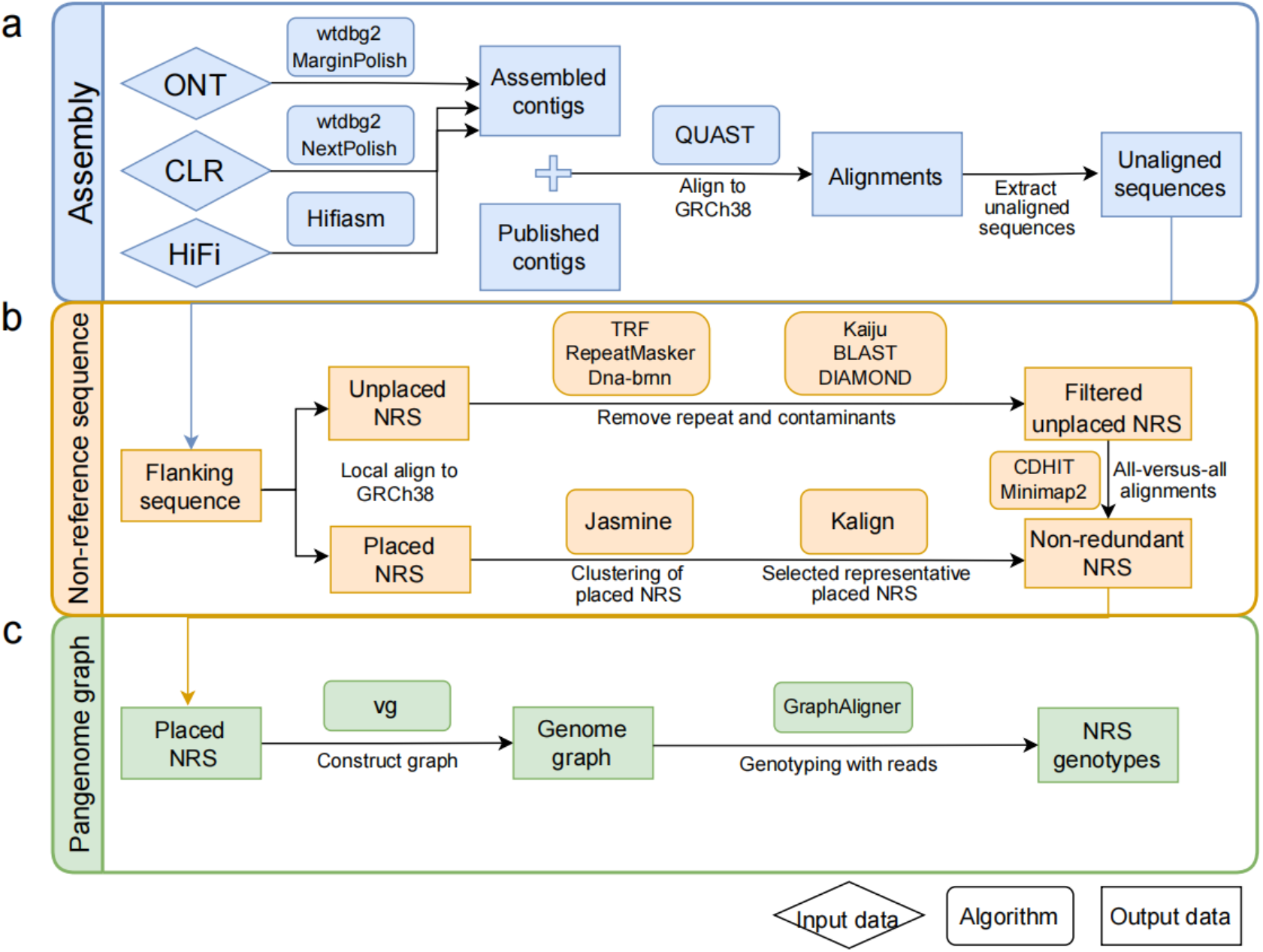
Schematic representation of GraphNRS. **a**, Long-read sequencing data from different platforms are *de novo* assembled and polished. **b**, The NRSs are anchored to GRCh38. Placed NRSs are clustered to select the representative NRSs, and unplaced NRSs are clustered after filtering out contaminants and centromeric repeats. Then, we merge the placed and the unplaced NRSs to obtain the nonredundant NRSs of the whole population. **c**, vg is used to construct the graph pangenome, and NRS genotyping is performed for each NRS of the individual.

To obtain an assembled high-contiguity genome, we first estimated the required sequencing depth of the long reads. We assembled the individual genome based on sequences randomly extracted from six ONT datasets. We found that the cumulative length of contigs assembled using 12-fold depth data was comparable to that of those assembled by 25-fold depth data (**Fig. S1a**). In addition, the N50 length of the assemblies was over 13 Mb with 15-fold depth data (**Fig. S1b**). To evaluate the accuracy of our assembly strategy, we applied it to the 15-fold HG002 datasets generated by ONT, PacBio CLR and HiFi, respectively, and compared the assemblies to the benchmark data described by Shumate et al.^14^. We found that the base-level error rates for ONT, CLR and HiFi were 0.93%, 0.49% and 0.12%, and the assembly disagreement counts were 152, 132 and 182, respectively (**Table S1**). The errors of the assemblies across the three platforms were lower or comparable to those of the assembly with more than 50-fold ONT reads (135 for assembly disagreements)^15^. The analysis suggested that our assembly strategy could provide a reliable assembly using sequencing data with around 15-fold depth.

After quality assessment, we individually *de novo* assembled 473 genomes sequenced by LRS platforms from previous studies and public database, with an average sequencing depth of 19.6-fold (**Table S2**, **Methods**). To further improve the assembly quality, we polished the assemblies except for those from the PacBio HiFi datasets. Additionally, we downloaded 66 publicly available assembled genomes sequenced by LRS platforms with quality assessment. Finally, we obtained 539 assemblies, where 431, 39 and 69 were obtained from the ONT, CLR and HiFi platforms, respectively (**Table S2**). The average length of 539 assemblies was 2,826 Mb (**Fig. 2a**), which recovered 93.9% of GRCh38 and 93.1% of the protein-coding sequences assessed by QUAST (**Fig. S2a**). These assemblies produced high-contiguity contigs, with an average N50 length of 16.3 Mb (**Fig. S2b**), which was much longer than the previously published genomes HX1 (8.3 Mb)^10^ and NH (3.6 Mb)^16^ that were both sequenced by the CLR platform. In addition, the draft assembly had a high level of macrosynteny with the reference genome GRCh38, confirming the accuracy of our assemblies (**Fig. S3**). Consistent with a previous study, we also found that LRS-based assembly detected errors that occurred in GRCh38, which was mostly based on BAC sequences and thus might result in multiple gaps and errors in the regions of scaffold switch-points^17^. For instance, in the switch-point of two original BACs, RP4-783C10 and RP11-109P14, a 2.4 kilobase (kb) sequence was missed in GRCh38 (**Fig. S4a**), and the missing sequence could be recovered in the assemblies in our study (**Fig. S4b**).

**Figure 2.**
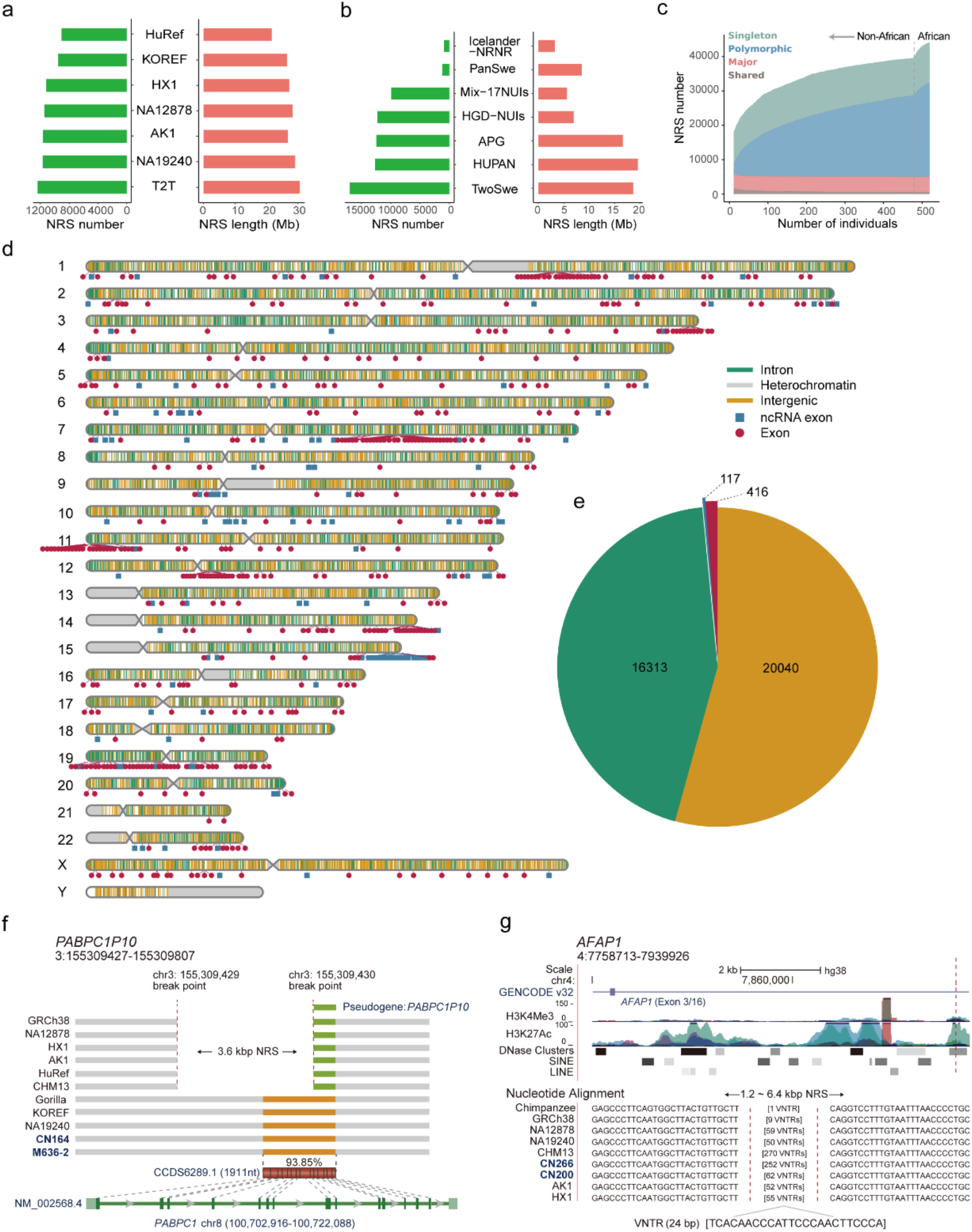
Characterization of NRSs for the whole population. **a**, Overlapped NRSs of this study to the different human genomes. **b**, Overlapped NRSs of this study to the different human pangenomes. **c**, The growth of the NRS number with an increase in sample size. The growth before the vertical dotted line is for non-Africans, and the growth after the vertical dotted line is for Africans. Four categories based on the AF are shown. **d**, Locations of the nonredundant NRSs against the reference genome GRCh38. **e**, No. of nonredundant NRSs intersected with different gene types. **f**, A 3.6 kb NRS anchored to the left end of pseudogene *PABPC1P10*. The green and orange bars represent *PABPC1P10* and the region with high identity to the CDS of *PABPC1*, respectively. The sample name in blue indicates the genome assembly generated in this study. **g**, An NRS composed of a VNTR with a 24-bp repeat unit present in nonhuman primate and multiple human genomes.

We extracted high-confidence NRSs from the assemblies using a hierarchical method (**Fig. 1 and Methods**). First, we aligned the assembled contigs to the reference genome GRCh38 and initially extracted an average length of 17.0 Mb raw NRSs. To obtain high-confidence NRSs, we removed contaminants, satellite sequences around the centromeric regions, contigs with ultralow or ultrahigh depths and unplaced singleton NRSs (**Methods**). To orthogonally evaluate the reliability of our identified NRSs, we assessed the quality of the NRSs of 10 samples in this study (**Table S3**) as well as HG002, which were sequenced by both ONT and PacBio HiFi^14^. We found that on average, 93.4% of NRSs (ranging from 92.0% to 94.8%) could be validated by PacBio HiFi data (**Table S3**), showing the reliability of the extracted NRSs. We then applied our method to the 539 genome assemblies. In total, we identified 5.1 million high-confidence NRSs for all the samples. And each individual obtained an average length of 6.3 Mb NRSs (**Fig. S5a**). After all the NRSs were merged and redundant NRSs were removed, we obtained 45,284 nonredundant NRSs with a cumulative length of 59.7 Mb, and the N50 length was 3.7 kb (**Fig. S5b**).

The *de novo* assembly strategy could identify NRSs both placed to GRCh38 or not. 36,853 (spanning 27.3 Mb) and 8,431 (32.4 Mb) NRSs were placed and unplaced relative to GRCh38, respectively (**Table 1** and **Table S4**). We compared the placed NRSs to the insertions (INSs) detected by the mapping strategy in a previous study^18^ and found that 11,150 (30.3%) NRSs were intersected with the INSs, suggesting that NRSs were reliable and our *de novo* assembly strategy revealed considerably more NRSs than the mapping strategy.

**Table 1.** Summary of the non-redundant NRSs

| Category | No. of<br>NRSs | Total length<br>(bp) | Average<br>Length (bp) | NRSs per<br>sample | Samples<br>per NRSs | No. of recovered<br>NRSs<sup>a</sup> | Length of recovered<br>NRSs (bp) |
| --- | --- | --- | --- | --- | --- | --- | --- |
| Placed | 36,853 | 27,259,120 | 740 | 8,577 | 125 | 11,511 | 7,669,944 |
| Unplaced | 8,431 | 32,409,688 | 3,844 | 834 | 53 | 3,814 | 27,696,129 |
| Total | 45,284 | 59,668,808 | 1,318 | 9,410 | 112 | 15,325 | 35,366,073 |

a: The number of non-redundant NRSs intersected with the previously published genomes and pangenomes (details in Methods).

The NRSs were compared to the previously published genomes and pangenomes^3^. Among the five human genomes we compared, the most overlapped NRSs were found in currently the most complete genome, T2T-CHM13^2^, including 12,377 NRSs with a total length of 30.7 Mb (**Fig. 2a**). For the pangenomes, HUPAN, consisting of NRSs from 275 Chinese individuals^6^, had the largest overlapping NRS length (19.6 Mb) owing to the highest proportion of the East Asian population in our study (**Fig. 2b**). In total, we confirmed 15,325 (33.8% of total) NRSs with a cumulative length of 35.4 (59.3%) Mb in the previous datasets (**Table 1**). The large number of novel NRSs (66.2%) showed that our study greatly expanded our current knowledge of the human genome.

### Distribution of NRS in the human genome

NRSs were nonrandomly distributed in the genome. For the 36,853 placed NRSs, 8,307 (22.5%) NRSs were in the last 5 Mb of chromosome arms (spanning 240 Mb), showing enrichment at the end of chromosome arms (odds ratio = 2.8, *P* = 8.8×10^−68^, Fisher’s exact test). In addition, we identified 144 hotspots spanning 141 Mb of the genome (**Table S5**). Of these, 112 (77.8%) hotspots were located in segmental duplications (SDs), which may be due to the increased mutation rate and interlocus gene conversion within SDs^19^.

For all the NRSs we identified, 78.5% were repeat sequences and 27.5% were nonrepeat sequences (**Table S6**). The percentage of repeat NRSs was in the range of some previous studies (75.0-88.6%)^6,11^. The repeat NRSs consisted of repeat elements, including variable number tandem repeats (VNTRs, 17.6%), short tandem repeats (STRs, 12.4%), short interspersed nuclear elements (SINEs, 11.0%) and long interspersed nuclear elements (LINEs, 14.8%) (**Table S6**). Previous studies showed that NRSs containing repeat elements had potential functional significance, such as producing variants underlying adaptive evolution^20^. The enrichment of repeat elements may be because these NRSs arise from the flanking locations with low-complexity sequences in genome where repetitive sequences are prone to expand.

### NRS in human populations and nonhuman primates

Among all the NRSs in the study, the shared NRSs with allele frequency (AF) of 1 had 1.1%, the major (1 > AF ≥ 0.5) had 10.9%, the polymorphic (0.5 > AF but not singleton) had 74.8% and the singleton (occurred in one sample) NRSs had 13.2%. It is noticeable that the low frequency NRSs accounted for the majority, with 69.4% at an AF less than 0.1 (**Fig. S6a**). Among all novel NRSs, 77.3% NRSs had an AF less than 0.1, showing the advantage of identifying low-frequency NRSs in a population study.

To understand how NRSs were distributed in regional populations in the world, we compared NRSs in the five major populations in the study, including East Asian (EAS), African (AFR), South Asian (SAS), European (EUR) and American (AMR). Out of the 45,284 NRSs, 38.7% (17,521) were common in all five populations, and 35.6% (16,112) were population specific (**Fig. S6b**). Previous studies showed that the African population had more SVs compared to non-Africans due to their large genetic diversity^17^. Here we also found that African had a significantly higher number of NRSs than non-African (*P* = 1.5 × 10^−30^, two-tailed *t* test, **Fig. S6c**).

To estimate how many individuals could capture the vast majority of NRSs in the population, we assessed the growth curve of the number of NRSs with respect to the population size. Because the African population had a much higher number of NRSs than non-African populations, we analysed the two groups separately. We observed that the number of the shared and the major NRSs stayed stable for the non-Africans and this trend did not change after adding Africans (**Fig. 2c**). In contrast, the number of polymorphic NRSs and the singletons gradually increased for the non-Africans and after adding Africans, these numbers quickly increased with a higher positive slope (**Fig. 2c**). This suggested that our study obtained the vast majority of the shared and the major NRSs for both non-Africans and Africans. And we need more samples to obtain the polymorphic NRSs and the singletons, particularly for Africans.

To understand the origin of NRSs, we compared the NRSs to the genomes of nonhuman primates, including chimpanzee^21^, gorilla^21^ and rhesus monkey^22^. Out of the 45,284 NRSs, 21.3% (9,635) were found in the genomes of nonhuman primates. The NRSs that overlapped with the human genome were consistent with primate divergence^23^, where 8,236, 6,538, and 2,393 NRSs were in the genomes of chimpanzee, gorilla, and rhesus monkey, respectively (**Fig. S6d**). In addition, 2,605 NRSs were specific to chimpanzee, suggesting that the NRSs were generated in the process of evolution and divergence. A total of 1,777 NRSs were present in all the three nonhuman primates, implying that these NRSs originated from a common great-ape ancestor (**Fig. S6d**).

### Functional annotation of NRS

Like other genetic variants, NRSs may be transcribed and have functional significance^24^. We first investigated whether NRSs contained known functional domains using the NCBI Conserved Domain Database (CDD)^25^ and the Pfam database^26^. We found that 118 annotated NRSs with known functional domains were associated with 134 genes. Of these, 119 (88.8%) genes can be validated by RNA sequencing data or the T2T-CHM13 genome (**Table S7**), indicating that these NRSs have the potential to be transcribed and have functional significance. All the 134 genes had homologous genes in nonhuman primates, suggesting that these genes may be derived from the gene duplication in evolution.

To further understand how NRSs interact with genes, particularly those relevant to disease, we annotated their inserted location. A total of 20,040 (54.4%) were in intergenic regions, and 16,813 (45.6%) overlapped with known genes (**Fig. 2d** and **Table S8**). For those in genic regions, 16,313 NRSs were in the introns of genes. 448 NRSs intersected with the exons of 322 protein-coding genes while 117 NRSs intersected with that of 96 non-coding genes (**Fig. 2e**). Notably, 77.0% of the NRSs that intersected with the exons of protein-coding genes tended to have a low AF (AF < 0.1) (**Fig. S6e**), indicating that these NRSs may be under negative selection^18^. Particularly, 84 NRSs intersected with exons of 71 protein-coding genes that are in the Online Mendelian Inheritance in Man (OMIM) catalogue (**Table S8**), suggesting that these NRSs may have a functional impact on disease.

STRs, consisting of tandemly repeated short (1 to 6 bp) DNA sequence motifs, were reported to cause more than 60 phenotypes^27^. Particularly, triplet repeats have been recently reported to be associated with multiple neurodegenerative disorders^28^. We identified 210 NRSs with triplet repeats, of which 102 NRSs intersected with 100 distinct genes, including 28 genes reported in the OMIM catalogue (**Table S9**). For instance, we detected a 678-copy CTG repeat expansion in *ATXN8OS*, which was reported to be associated with amyotrophic lateral sclerosis^29^; a 429-copy gain of a CGG repeat in *ZNF713*, which was reported to be associated with the folate-sensitive fragile site FRA7A^30^; and a 235-copy gain of ACC repeats in *GRIK4*, which contributed to the risk of schizophrenia^31^.

VNTRs are tandem repeats with motif length ≥7 bp and have been reported to affect diverse human phenotypes through intersecting with protein-coding exons^32^. We found 19 NRSs composed of VNTRs located in the exons of the protein-coding genes, including nine reported in a previous study (**Table S10**)^32^. We observed that several VNTRs located in mucin family genes, such as *MUC2* and *MUC6*. Previous studies showed the association between *MUC6* VNTR expansion and Alzheimer pathologic severity^33^. In this study, we detected the expansion of VNTR located in *MUC6* with 672-bp repeats (copies ranging from 2.5 to 7.1). Similarly, in *MUC2*, we detected variable expansions with 24-bp repeats (copies ranging from 37.5 to 141.9). Our datasets provide a resource to investigate the contribution of tandem repeats expansion to phenotypes.

Annotated NRSs could also provide important clues into gene evolution. For instance, a 3.6-kb NRS inserted into the front of a truncated pseudogene *PABPC1P10* (**Fig. 2f**), which is homologous to *PABPC1*, functions in regulating the metabolism of mRNA^34^. This indicates that the sequence containing the complete pseudogene for *PABPC1* existed before human divergence, suggesting that the pseudogene might be redundant or that loss-of-function (LoF) variants were tolerated during human evolution. In contrast, some NRSs expanded after human divergence. We found a novel NRS composed of a VNTR with a repeat unit of 24 bp (**Fig. 2g**) in the second intron of *AFAP1*, which was associated with small intestine cancer and open-angle glaucoma^35^. This NRS also intersected with H3K27Ac, H3K4Me3 and DNase clusters, indicating that it might play a role in regulating gene expression. We found one copy of this VNTR in the chimpanzee genome and 50–270 copies in the genomes of diverse human populations. This finding suggested that the VNTR in *AFAP1* expanded after human divergence from a common great ape ancestor and was variably present in extant individuals.

### Construction and utility of a graph-based pangenome of human NRS

The linear reference genome is known to be inadequate and biased to detect genetic variations^7^. Recent studies have shown that the graph genome can effectively genotype variants due to the higher sensitivity of the graph-based alignment approach than the linear reference-based alignment approach^36,37^. To build a graph-based genome, we used the current reference genome GRCh38 as the backbone and constructed a graph genome using the 36,853 placed NRSs as the nodes, which can generate alternative paths (**Fig. 1c** and **Methods**).

To test whether the graph genome of NRS improves the alignment, we mapped the public DNA and RNA short-read sequences from diverse populations to GRCh38 and our pangenome. We observed that the mapping rate was increased for both the DNA sequences (from 97.69% to 99.28%, *P* = 0.0059, Wilcoxon signed-rank test) and the RNA sequences (from 97.43% to 98.50%, *P* = 0.002, Wilcoxon signed-rank test) (**Fig. S7**). This result showed that the graph pangenome could improve the detection of novel variants and the quantification of gene expression.

To assess the genotyping accuracy from the graph genome, we first compared the genotypes of NRSs that overlapped with INSs of HG002. The sensitivity of the genotypes of NRSs was 0.94 (**Table S11**), which was higher than that of the non-SNPs (0.89 in 1000 Genome + GIAB) reported in a previous study^38^, while the precision was 0.96. We further validated the genotypes using the public trio datasets^39^. For the genotypes of the parents of six trios, the average abnormal proportions of offspring with Mendelian inheritance ranged from 0.35% to 7.73%, with an average of 2.13%, showing lower genotyping error (**Table S12**). To further assess the genotyping performance at the population scale, we analysed the AF and heterozygosity of the NRSs and examined how many NRSs fit Hardy-Weinberg equilibrium (HWE) (**Fig. S8**). We found that 94.0% of the NRSs showed no significant deviation when testing for HWE, which was higher than a previous study for SVs (HWE = 90.7 - 90.9%)^40^. These analyses suggested that our graph genome can reliably genotype the NRSs.

### NRS improves eQTL detection

It has been suggested that NRSs could be the causal variant for eQTLs, and their longer length is more likely to alter the gene expression compared with SNPs^41^. We conducted eQTL analysis using the graph pangenome. We integrated the NRS genotypes and RNA-seq data to assess eQTLs for 451 samples from the Genetic European Variation in Disease (GEUVADIS) consortium. Principal component analysis (PCA) based on the graph-based genotyping of NRSs showed that the samples consisted of four European ancestry and one African ancestry populations^42^, consistent with the PCA result from GEUVADIS based on SNPs (**Fig. S9a**). We next performed association analysis between the transcript expression levels and the graph-based genotypes of NRSs within 1 Mb from the transcription start site (TSS)^43^. Finally, we identified a total of 565 NRS-transcript pairs with significant expression associations with a false discovery rate (FDR) less than 0.05 (**Fig. 3a** and **3b**). Of these, 98 eQTLs were previously reported and 467 (82.7%) were novel^17,43^ (**Table S13**).

**Figure 3.**
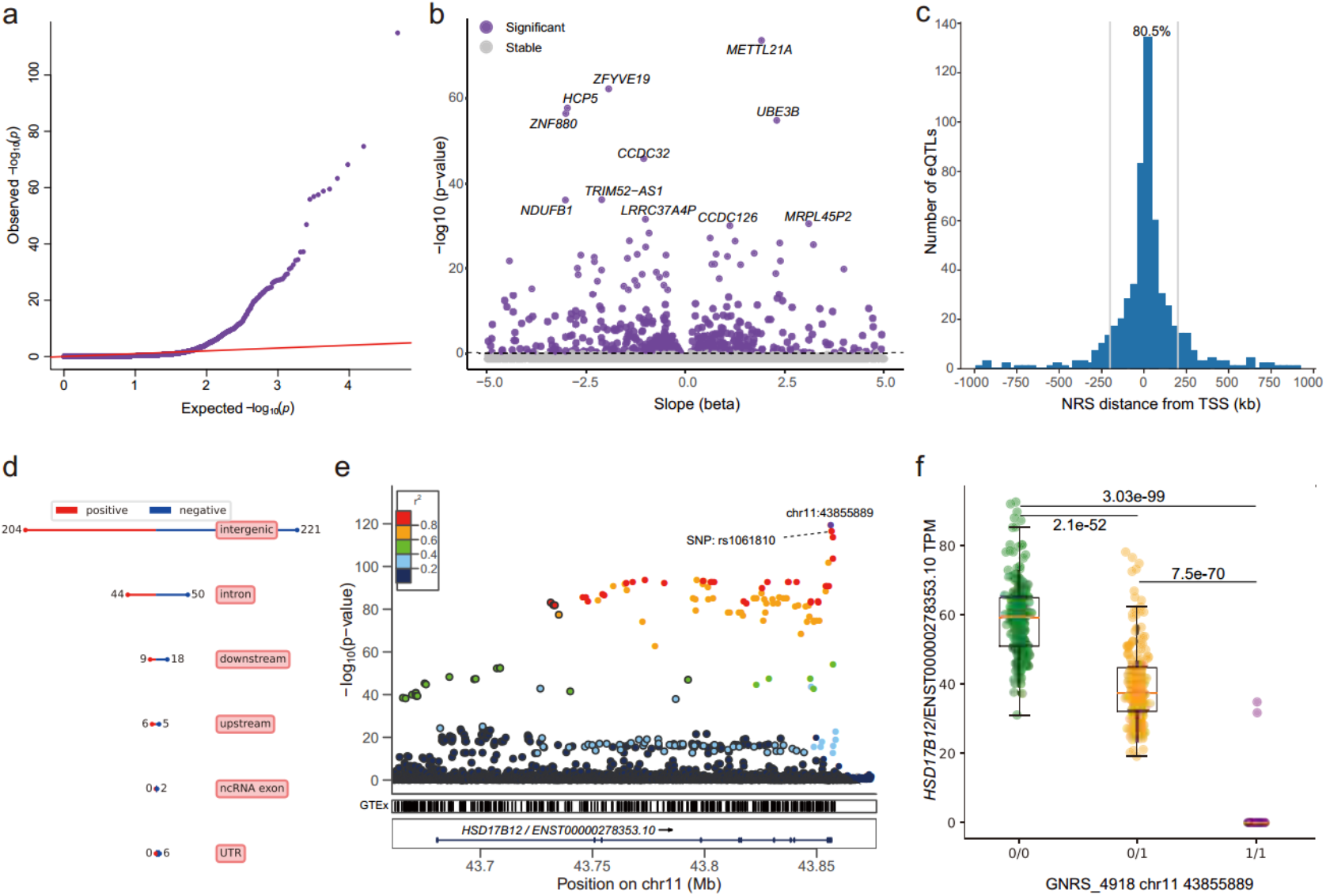
eQTL analysis based on the graph pangenome. **a**, Quantile-quantile plot of permutation p values for all NRS transcript pairs tested. **b**, Volcano plot of eQTLs and the estimated effect (beta) of the alternative NRS allele on transcript expression. **c**, Distribution of the distance of significant NRS eQTLs from the TSS of the associated genes. The blue line indicates the position of 200 kb from the TSS. **d**, Summary of the annotation and impact of eQTL-associated NRSs. “positive” and “negative” indicate the NRS allele increases and decreases the expression of corresponding transcript. **e**, An example of NRS lead-eQTL for *HSD17B12*. The y-axis represents the significance of the association, with the top eQTL being the highest point. The colours indicate the LD values between the top signal and other variants. **f**, The median transcripts per million (TPMs) for the transcript (ENST00000278353) of *HSD17B12* for individuals containing different NRS alleles. “0/0”: homozygous allele same as the reference (n = 187); “0/1”: heterozygous NRS (n = 211); “1/1”: homozygous NRS (n = 53). Boxes represent the median and quartiles, whiskers extend from the box up to 1.5 times the interquartile range. The p values between different alleles were calculated based on a two-tailed t test.

NRSs with significant effects on gene expression tended to be near the genes that they regulated, with 80.5% of significant NRS eQTLs occurring within 200 kb of the corresponding TSS (**Fig. 3c**), which was consistent with a previous study^43^. The NRSs could both upregulate (263 NRSs) and downregulate (302 NRSs) the expression of nearby genes. However, NRSs tended to downregulate the gene expression for those located in the exon region and downstream of genes (**Fig. 3d**). Eight NRSs were in exons and had a significant effect on the corresponding genes (**Table S14**). In addition, NRSs may have different impacts on distinct transcripts of the same gene, which may be related to the exact location of the NRS. Among the NRS-gene pairs, 425 unique genes were significantly enriched in MHC class II receptor activity (GO:0032395) in the gene ontology (GO) analysis (odds ratio = 46.6, adjusted *P* = 3.5×10^−4^, Fisher’s exact test, **Table S15**), suggesting that NRSs might mediate immune diversity by regulating transcript levels. Furthermore, to understand how NRSs affect the expression of genes, we annotated these NRSs using epigenetic state information generated by the Roadmap Epigenetics Consortium (REC)^44^. The NRSs were intersected with 12 states (**Table S13**) and strongly enriched in the transcribed state at the 5’ and 3’ end of genes (odds ratio = 17.0, adjusted *P* = 1.6×10^−6^, Fisher’s exact test, **Fig. S10**), as well as the states of TSS, transcription and enhancers. In addition, we found a strong depletion of expression associations for NRSs that intersected with the state of quiescent that is devoid of important epigenetic marks (odds ratio = 0.3, adjusted *P* = 1.4×10^−23^, Fisher’s exact test, **Fig. S10**). It suggested that these NRSs might interfere the epigenetic status and thereby affect the gene expression.

Compared to SNPs for eQTL analysis, NRSs could improve the power of detecting eQTLs. We observed that 15 NRSs more significantly affected gene expression than SNPs, and six of them were in the genic region of the affected genes (**Table S16**). For instance, many SNPs were found to be significantly associated with the transcript expression of *HSD17B12* (ENST00000278353), which encodes 17 beta-hydroxysteroid dehydrogenase and is associated with long-chain fatty acid metabolism^45^. The top signal of SNPs, rs1061810 (*P* = 2.3×10^−117^), was in the 3’UTR of the transcript. Interestingly, we observed that an NRS (GNRS_4918, chr11:43855889, 318 bp), which was also located in the 3’UTR and was in high LD (r^2^ = 0.87) with the SNP rs1061810, which was even more significant than the top SNP signal (*P* = 5.6×10^−120^) (**Fig. 3e**). Three genotypes of this NRS in the population were in accordance with HWE (*P* = 0.61), indicating accurate genotyping. Furthermore, the median transcripts per million (TPMs) of this transcript with genotypes “0/0”, “0/1” and “1/1” were 59.2, 39.6 and 1.3, respectively, suggesting that the NRS negatively regulated the transcript level (**Fig. 3f**). Although both the SNP and the NRS are in the 3’UTR of the transcript, the NRS has a longer length and is prone to directly regulate the transcript level of this gene^41^. This demonstrated that our graph genome could improve eQTL detection to identify NRSs that may be the causal variant candidate of the differential transcript expressions.

### NRS contributes to the local adaptation of diverse populations

To determine NRSs that underwent population adaptation, we assessed population differentiation using the population branch statistic (PBS) based on the genotypes of NRSs for East Asian, African, and American populations, which had large proportions of individuals in our study. We detected 39 unique NRSs with significant PBS scores (99.9% rank) located in or flanking 38 unique genes, which indicated the potential adaptation loci associated with different populations (**Fig. 4**, **Tables S17 and S18**). Six genes, namely, *KCNH7*, *LARS2*, *LUZP2*, *SLC30A9*, *SLC37A1* and *TAF1B*, were reported by the catalogue of human genome adaptation^46^, verifying the ability of NRSs to reveal signals of local adaptation.

**Figure 4.**
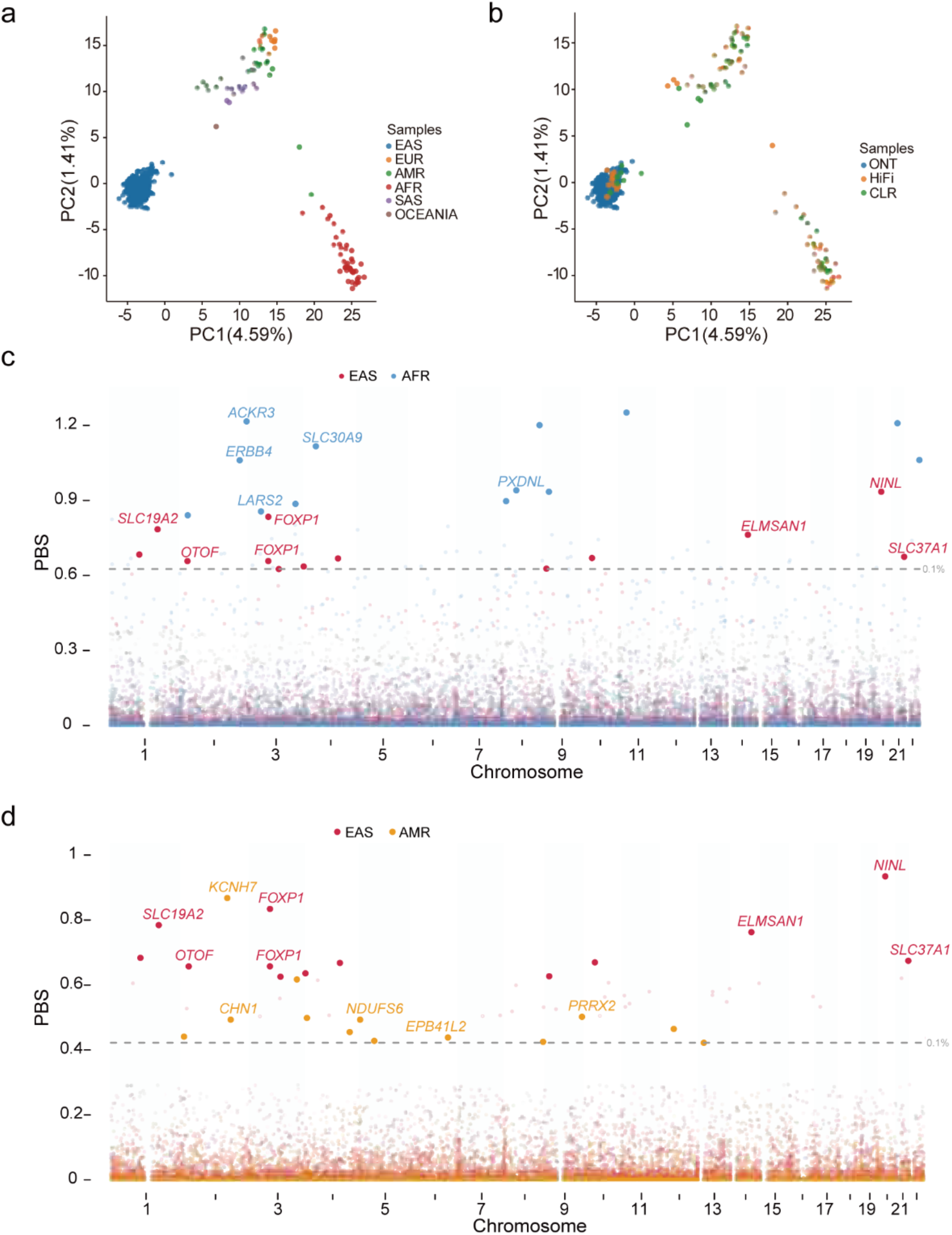
Local adaptation of the NRSs in diverse populations. **a**, PCA of all the samples across different populations. The values in parentheses indicate the genetic variations explained by the first two PCs. **b**, PCA of all the samples across different sequencing platforms. **c**, PBS of NRSs for East Asians and Africans. The grey dotted line represents the top 0.1% (0.62) of the PBS ranked score. **d**, PBS of NRSs for East Asians and Americans. The grey dotted line represents the top 0.1% (0.42) of the PBS ranked score.

We found that two NRSs located in the intron of *FOXP1* with large frequency differences between East Asian population and African/American populations. For instance, the frequencies of a 317-bp NRS (GNRS_20923) were respectively 0.72 and 0.54 in Africans and Americans, while the frequency was as low as 0.04 in East Asians (**Fig. 4**, **Tables S17 and S18**). Previous study showed that *FOXP1*, encoding a member of the forkhead box (FOX) transcription factor family, may play a role in human speech and language phenotypes^47^. The languages of East Asia belong to several distinct language families, of which Chinese is the language with an idiographic nature, compared to other languages with an alphabetic nature^48^, and influences the neighbouring Asian languages due to the dominance of Chinese culture in East Asia. A study of adult mouse single-cell RNA reported that *FOXP1* was expressed in the cortex^49^, and functional magnetic resonance imaging showed neural signals in different parts of the temporal and frontal cortex for different languages^50^, suggesting the potential function of *FOXP1* in regulating different language systems. The differentiation of this NRS in *FOXP1* may be the result of local adaptation for long-term mastery of different language systems.

Some NRSs were significantly associated with genes related to metabolism, such as three members of the solute carrier (SLC) superfamily (**Fig. 4**, **Tables S17 and S18**), which is responsible for transporting extraordinarily diverse solutes across biological membranes. For example, we observed an NRS (319 bp) in the intron of *SLC30A9* encoding a zinc transporter, with AFs of 0.96 and 0.12 in the East Asian and African populations, respectively. It has been suggested in a previous study that *SLC30A9* was under natural selection in both East Asians and Africans but in different directions, which is the result of local adaptation of human populations due to the local zinc state or diets^51^. We also detected a 654-bp NRS in the intron of *SLC37A1*. The gene is associated with glucose homeostasis and sugar transport^52^. It has also been shown to encode the transmembrane protein involved in ion transport and is therefore likely to influence milk mineral composition^53^. People in most parts of Europe and some parts of Africa underwent convergent adaptation of lactase persistence to improve the ability to digest milk^54^. The lactose intolerance of Asians is mainly due to the lack of the lactase enzyme with the function of breaking down lactose into smaller sugars called glucose and galactose^55^ and may subsequently lead to the adaptation of *SLC37A1* accompanied by adaptation to break down lactose efficiently. Additionally, an NRS (654 bp) was found to be in the intron of *SLC19A2*, which was reported to be associated with diabetes mellitus and thiamine-responsive megaloblastic anaemia^56,57^. The gene was confirmed to be under selection in the Asian population^58^.

In addition, we found NRSs with adaptation signals associated with anaemia. For instance, a 320-bp NRS located in the intron of *LARS2* (**Table S17**), which encodes a mitochondrial aminoacyl-tRNA synthetase, was reported to be associated with sideroblastic anaemia (MIM: 617021)^59^ and red blood cell count^60^. A 330-bp NRS was 501 kb downstream of *LUZP2*, which encodes a leucine zipper protein and was reported to be associated with serum iron levels^61^. Indeed, a previous study reported significant natural selection signatures covering *LUZP2*^62^ and *LARS2*^46^ in Japanese and Central/South Asians (STU), respectively. Iron deficiency is the most common cause of anaemia worldwide^63^. The age-standardized rate of anaemia in Africa (ranging from 24.6% to 44.2%) was higher than that in East Asia (20.5%). In particular, the highest national prevalence of anaemia (95.0%) was found in Zambia^64^. The severe anaemia status in Africa may be due to both environmental factors, such as economic underdevelopment and nutritional deficiencies, and genetic factors, where special alleles of genes are associated with the iron regulation of serum.

Furthermore, we detected NRSs with adaptation signals associated with type 2 diabetes (T2D). An NRS (332 bp) located 23.8 kb downstream of *PRDM5* (**Table S17**), a gene involved in fasting plasma glucose in a GWAS with diabetes-related traits^65^, was also reported as a signature of geographically restricted adaptation in all three populations of Africans^66^. We also found an NRS (342 bp) located in the intron of *ERBB4*, which is a member of the Tyr protein kinase family and the epidermal growth factor receptor subfamily. Genetic studies have indicated a link between *ERBB4* and T2D and obesity^67^. Furthermore, an NRS (349 bp) was found to be 9.6 kb upstream of *PIM3* and intersect with H3K27Ac, H3K4Me1 and transcription factor (TF) clusters (**Fig. S11a**), which belong to the Ser/Thr protein kinase family. This gene was reported to be associated with T2D by aggregating genome-wide genotyping data from 32 European-descent GWASs (*P* = 2.0 × 10^−8^)^68^. In addition, an NRS (343 bp) was located 158 kb upstream of *DMRTA1*. The SNP (rs1575972) near *DMRTA1* was reported to be significantly associated with T2D (*P* = 4.7×10^−13^)^69^. The NRS was in the region flanking three SNPs that were significantly associated with diabetes (*P* = 2.4×10^−11^ to 1.8×10^−11^) and in high LD (r^2^ > 0.8) with the top signal (**Fig. S11b**). Asians have been shown to have a higher prevalence of diabetes than Africans^70^. Our analysis suggested that the adaptation of the NRSs related to these diabetes-related genes might contribute to the difference in diabetes incidence in different populations.

### Phenotypic association of NRS

We next conducted GWAS on NRS genotypes and clinical traits using our constructed graph genome. Our study included 327 samples with 68 traits obtained during health check-ups^18^. We fitted an additive genetic model with relevant covariates for the quantitative traits using the 5,643 NRSs with a minor allele frequency (MAF) of > 0.05. The genomic inflation factor (λGC) ranged from 1.00 to 1.05, with an average of 1.01, suggesting very low inflation. Finally, we identified 14 NRSs significantly associated with eight phenotypes (*P* < 8.9×10^−6^, the Bonferroni-corrected significance threshold, **Fig. 5a** and **Table S19**). For example, one of these NRSs, GNRS_28218, was significantly associated with mean corpuscular haemoglobin (MCH), mean corpuscular volume (MCV) and red cell volume distribution width-coefficient of variation (RDW-CV), which are indicators for assessing anaemia. GNRS_28218 is in the intron of the interaction protein for cytohesin exchange factors 1 (*IPCEF1*). A previous GWAS of chronic lymphocytic leukaemia (CLL) identified a susceptibility locus mapping to *IPCEF1* (rs2236256, *P* = 2×10^−10^)^71^. CLL is frequently complicated by cytopaenias, either due to bone marrow infiltration or autoimmunity, and results in autoimmune haemolytic anaemia (AIHA)^72^, suggesting the potential contribution of *IPCEF1* to anaemia. GO annotation shows that this gene is related to peroxidase activity (GO:0004601) and oxygen carrier activity (GO:0005344). The GenomeRNAi database shows that the human phenotype of RNA interference with *IPCEF1* is decreased endocytosis of transferrin^73^, which has an impact on iron incorporation by erythroblasts. The above evidence suggested that GNRS_28218 associated with *IPCEF1* was likely to have a functional impact on anaemia.

**Figure 5.**
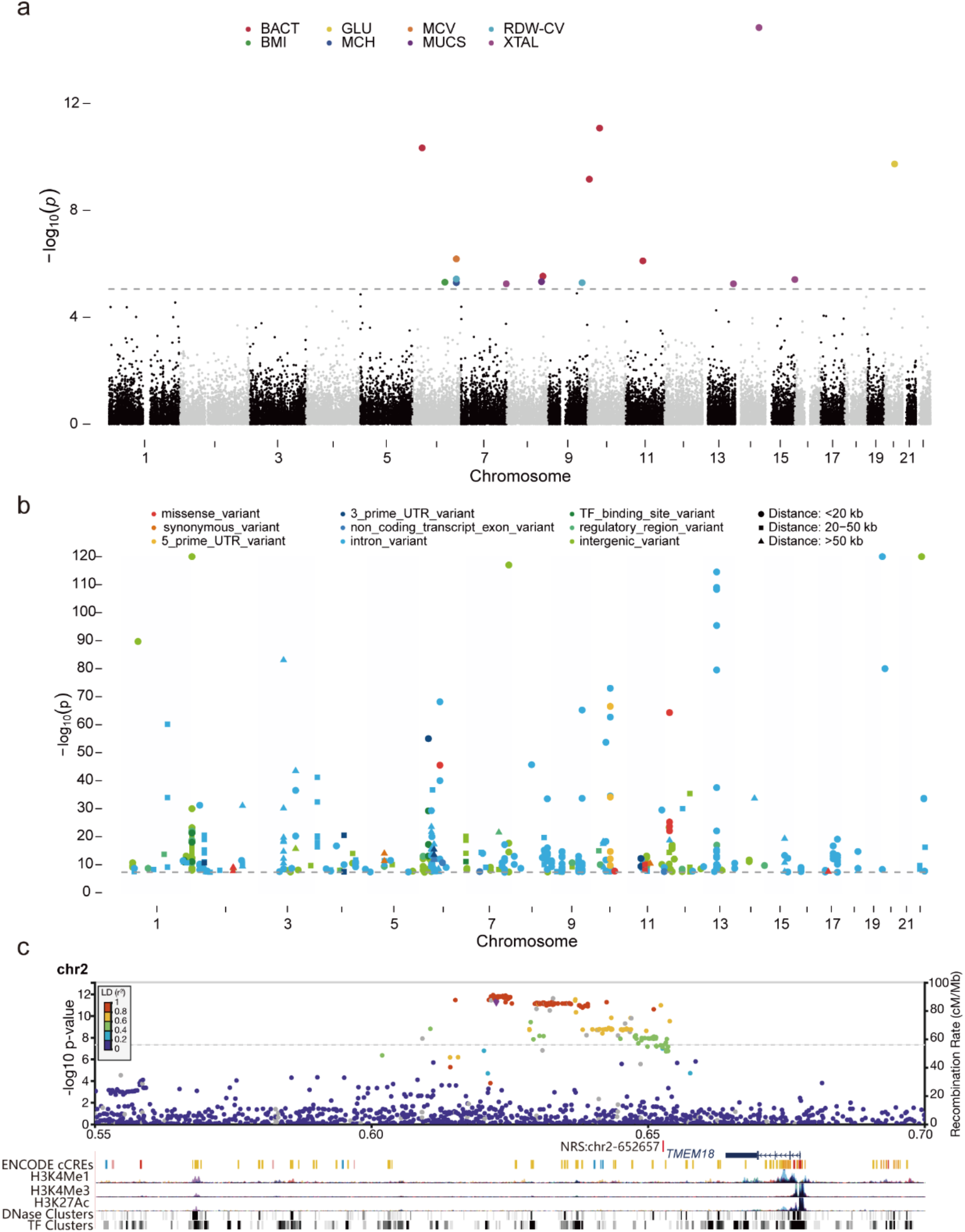
NRSs. significantly associated with phenotypes. **a**, Manhattan plots show NRSs plotted on the x-axis according to their position on each chromosome against, on the y-axis (shown as −log10 P value), the association with clinical phenotypes. The grey dotted line indicates the significance threshold (p = 8.9×10^−6^ through Bonferroni correction). BACT: urinary bacteria, GLU: blood glucose, MCV: mean corpuscular volume, RDW-CV: red cell volume distribution width-coefficient of variation, BMI: body mass index, MCH: mean corpuscular haemoglobin, MUCS: mucus, XTAL: urinary crystal. **b**, Manhattan plot for the phenotype-associated SNPs that are in strong LD (r^2^ > 0.8) with NRSs. Different colours demonstrate different gene features in the GWAS catalogue, and the shapes indicate the distances between the SNP and the NRS. **c**, Regional SNP association plots with the NRS (red vertical line) around *TMEM18* shown in high LD (r^2^ > 0.8) with the top signal of BMI.

To further explore the potential functions of NRSs in phenotypes, we identified NRSs that were in LD with SNPs associated with phenotypes in the GWAS catalogue^74^. Using a window size of 100 kb, we observed strong LD (r^2^ > 0.8) between 154 NRSs and 258 phenotype-associated SNPs at genome-wide significance (*P* = 5×10^−8^) reported in the GWAS catalogue (**Fig. 5b**, **Table S20** and **Methods**). We found that an NRS, GNRS_15339 on chromosome 2 (2p25.3), was 266 bp away from cis-regulatory elements and in strong LD with 15 unique SNPs, which were reported in regulatory or intergenic regions near *TMEM18* in 15 GWASs and were significantly associated with body mass index (BMI) (**Fig. 5c**). We found that seven additional NRSs (GNRS_21065, 27160, 30214, 30546, 33543, 12548 and 12873) were in strong LD with SNPs that were also significantly associated with BMI. All these SNPs were in the intergenic, regulatory or intron regions except for a missense variant (rs17826219, *P* = 3×10^−8^) in *ATAD5*. However, we noticed that the above eight NRSs detected in this study were not significantly associated with BMI (*P* = 0.0046 for GNRS_30214) in our GWAS, suggesting that a larger population size is needed in future study. Interestingly, among the 258 phenotype-associated SNPs that are in strong LD with NRSs, only 11 (4.3%) SNPs were in the coding region, of which seven and four SNPs were missense variants and synonymous variants, respectively. Furthermore, 25 (16.2%) NRSs with high LD with these phenotype-associated SNPs were validated to significantly regulate the gene expression by the eQTL analysis (**Table S20**). Considering that most significant NRSs and SNPs were not located in the coding region of genes but NRSs extended further (in terms of bases) than SNPs and were more likely to affect the nearby gene expression, the NRSs that presented strong LD with these significant SNPs are more likely the causal variants to the corresponding phenotypes.

## Discussion

Many genetic sequences, particularly in the human populations, are missing from the current reference genome. Construction of a human pangenome reference that captures genetic variation in the population is critical in understanding genetic instructions in human^3^. Aiming to better characterize human NRSs and understand their functional significance, we conducted an LRS-based NRS study for 539 human genomes. Our method of identifying NRSs from the assemblies of LRS data showed a clear advantage because the majority of NRSs consisted of low-complexity sequences. The average length of NRSs extracted for each individual in this study (17.0 Mb) was slightly longer than the length in previous long-read assemblies (from 12.8 to 16.0 Mb)^3^, and was much longer than the length of NRSs from short reads (from 0.2 to 2.5 Mb). More importantly, our LRS-based datasets included NRS originating from multiple populations and provided a more complete pangenome than some previous studies mainly focusing on single population or with a small sample size^5,75,76^. As a result, of all the NRSs we identified, 66.2% (29,959) were novel. A total of 118 NRSs were annotated with known functional domains and were associated with 134 genes, of which 88.8% were validated by the RNA dataset or T2T-CHM13. 448 NRSs were in the exons of 322 protein-coding genes, indicating their potential to disrupt gene function. In addition, we showed that distribution of NRS on the human genome and in the human populations, as well as non-human primates, which gave us a better understanding of how NRS evolves. Our analysis extended our knowledge about how widespread the human NRS is and its functional impact on evolution, phenotypes and diseases.

We showed that the mapping rate of sequences was improved by applying a graph pangenome of NRSs, which has the potential to discover more variants and thus provides us with more complete genetic information for phenotypes and diseases. More importantly, the graph pangenome enables us to genotype NRS, which provides us with an opportunity to discover novel associations between NRS and phenotypes. We showed that NRS could improve the power of detecting eQTLs compared to SNPs. The novel eQTLs provided clues about the gene expression regulation by NRSs. Furthermore, many genotyped NRSs were found to be in strong LD with phenotype-associated SNPs in the GWAS catalogue. Most of these SNPs (95.7%) were in noncoding regions of genes, and therefore, very few of them were causal variants. These findings confirm that a vast number of genetic variants have not been discovered, and these undiscovered variants may be the missing causal variants. Similarly, the recently assembled complete genome T2T-CHM13 revealed hundreds of thousands of previously unresolved variants and the existence of additional copies of medically relevant genes when previous sequencing datasets of diverse populations were reanalysed^77^, further illustrating the importance of NRSs in genetic and genome research. Therefore, the graph pangenome of NRSs constructed in this study provides an import resource to reanalyse the relationship between these missing genetic variants and phenotypes and to narrow down the candidate causal variants or genes to phenotypes and diseases.

Currently, many NRSs cannot be anchored to the reference genome, mainly due to the breakage of the assembly in larger repeat regions. Recent advances in combining ONT ultralong and PacBio HiFi reads show promise in detecting the NRSs of repeat regions, particularly SDs and centromeric regions, by performing T2T assembly for the haploid CHM13^2^. And by adding deep sequencing data across more multiple platforms, such as Bionano optical maps and Hi-C Illumina short-read sequencing, Liao et al. obtained 47 state-of-the-art phased, diploid assemblies in HPRC^13^. Therefore, more deep sequencing data across multiple platforms are needed in future studies to assemble high-quality phased genomes and accurately identify NRSs in low-complexity regions such as SDs. In addition, to increase the power for the analyses of local adaptation and GWAS, larger sample sizes coupled with more phenotypes are needed for the diverse populations, particularly for the African population, due to its higher genetic diversity than other populations. As the sample size and genetic diversity increase, the number of nodes with multiple variants will increase in the graph, which will further improve the accuracy of alignment and genotyping.

In summary, our efforts of identification and characterization of human NRS provide a valuable resource to the human genomic research. The constructed graph-based pangenome also provides us with the opportunity to conduct eQTL, population local adaptation and genotype-phenotype association analyses, which will be an important step towards a global human pangenome.

## Methods

### Samples and datasets

In this study, 539 genomes sequenced by whole-genome LRS were *de novo* assembled, including 405 Chinese individuals sequenced by the ONT platform^18^ and 134 individuals from diverse populations sequenced by the ONT, PacBio CLR and PacBio HiFi platforms. Of which, assemblies and corresponding sequences of 65 individuals were directly downloaded, including 47 high-quality phased, diploid assemblies from HPRC^13^ (**Table S2**). To obtain high quality genome assembly, all the LRS data were under quality control by trimming 30 bases of start and 20 bases of end for raw reads and kept the reads with length longer than 500 bp^18^.

### *De novo* genome assembly of LRS datasets

To explore the relationship between the *de novo* assembly continuity and sequencing depth, we randomly selected reads with depths of 2×, 4×, 8×, 12×, 15×, 18× and 22× from six samples with sequencing depth larger than 25-fold. The down-sampling reads with different depths were then used for *de novo* assembly by wtdbg2^78^ v2.5 with parameters “ -p 19 -AS 2 -s 0.05 -L 500”. For *de novo* assembly with ONT data, we applied wtdbg2 using parameters mentioned above. To improve base accuracy of the assembly, the assembled contigs were polished by MarginPolish^15^ v1.3.0. The downloaded PacBio CLR and PacBio HiFi reads were assembled using wtdbg2 and hifiasm^79^ v0.16.1-r375 with default parameters, respectively. And assemblies from PacBio CLR reads were polished using NextPolish (https://github.com/Nextomics/NextPolish) v1.4.0 with parameters “-r clr -sp”.

We evaluated the completeness of assemblies and protein coding genes using QUAST^80^ v5.0.2 after aligning assembled contigs to primary assembly genome GRCh38 without ALTs^10^ coupled with corresponding gff annotation file (v95). To further assess base-level error rate and assembly disagreement counts, we applied the estimate methods used by Shafin et al.^15^. We randomly selected 15-fold sequencing data across the ONT, PacBio CLR and HiFi platforms for HG002 and independently assembled the genomes using the strategy in this study. We compared the assemblies to the benchmark data described by HG002 v1.7, which had high-quality genome based on datasets from multiple platforms^14^.

### NRS Detection

We applied a hierarchical strategy to extract NRSs (**Fig. 1a**). For each individual, the NRSs were generated from unaligned contigs by QUAST using commond “quast --no-gc --no-plots --no-html --no-snps --min-contig 1000 -o output -r ref_genome.fa -g ref_genome.gff -t threads assembly_genome.fa”^6^. To further confirm the unaligned sequences absent from the reference genome, the reference GRCh38.p13, which contained patches scaffolds, alternate loci and mitochondrial sequence, coupled with Epstein-Barr virus sequences (AJ507799.2) and decoy sequences (GCA_000786075.2), was used for further alignment. Then we aligned the clean LRS reads to the assembled contigs, and estimated the depths using mosdepth v0.2.5^81^. We filtered NRSs which were defined as collapsed or low read depth if these depths were greater than three folds or lower than one third of mean depth corresponding individual.

It is known that heterochromatic and centromeric regions are composed of tandem repeats HSat2,3 and Alpha satellites, which contributed to most of gaps of genome assembly^82,83^. RepeatMasker (http://www.repeatmasker.org) v4.0.9 and dna-brnn^84^ v0.1-r65 were used to specifically identify and remove Alpha and HSat2,3 if masked length ≥ 80% of total length.

Mosè Manni et al. reported that NRSs were overestimated in some previously published human pan-genome studies because most of them were contaminated with bacteria associated with original samples or introduced later during sequencing experiments, though some works had been done for filtering out contaminants in those studies. In this study, we used the method recommended by Mosè Manni et al.^85^ to remove contaminants (**Fig. 1b**). First, to avoid spurious matches derived from low-complexity and repetitive regions, we masked these low-complexity sequences with RepeatMasker and TRF^86^ v4.09 based on Dfam^87^ v3.0 and RepBase^88^ (v2018-10-26). Next, the remaining unaligned sequences were classified using Kaiju^89^ v1.7.3 with parameters “-t nodes.dmp -f kaiju_db_nr_euk.fmi -i non_ref.fa -a mem -z threads -o kaiju.out –v”, which had a good recall rate to detect more divergent sequences at amino acid level^85^. Through aligning above sequences against the pre-formatted “nr + euk” database (v2019-06-25), which contained protein sequences from bacteria, archaea, viruses, fungi, and microbial eukaryotes, we obtained a taxonomic classification of each unaligned sequences based on the continuous alignment with at least 100 amino acids and marked the sequences with label of “non-chordate”. Then, we searched the sequences against the nr and nt (v2020-1-9) databases using DIAMOND^90^ v0.9.21.122 and BLAST^91^ v2.10.0+, respectively. Based on the alignment and taxonomic classification, we retrieved the sequences labelled with chordate which were originally labelled as non-chordate by Kaiju. To get reliable NRSs, we considered any sequences labelled as non-chordates to be contaminants and removed them from further analysis.

To assess the effect of population size on the non-redundant NRS number, we randomly down sampled the individuals excluding the ones of PacBio CLR in this study. Due to the higher genetic diversity and more distinct SVs of Africans reported in the previous study^92^, we needed to classify all of the individuals into two categories, namely non-Africans and Africans. We added one individual for each step and repeated 10 times. And the NRS number was calculated based on the average of NRS number of down sampled individuals.

### Anchoring and validation of NRS

To anchor NRSs, we extracted the two flanking sequences with 1 kb length of the identified unaligned sequences and separately aligned them to the reference genome GRCh38 using AGE^93^ v0.4. If the alignment length of the flanking sequence was greater than 500 bp, and the coordinates of the two ends were less than 20 bp, the original sequence was considered to be successfully anchored to the reference genome. The locations of placed NRSs relative to reference genome were plotted using in-house script modified from RIdeogram^94^ v0.2.2. The sequences that still unaligned were considered as unplaced. If neither upstream nor downstream sequences could successfully align to the reference, the sequences were also regarded as unplaced. To further obtain reliable unplaced sequences, the unplaced sequences and contigs were realigned to genome using minimap2. Unaligned sequences were removed if aligned identity ≥ 90% and aligned length coverage ≥ 80% of total length. Furthermore, we filtered out unplaced sequences with a strategy reported in a previous study by the following conditions: (1) TRF reports > 80% masked bases; (2) RepeatMasker reports > 80% combined masked bases in satellites, simple repeat, and low complexity sequence categories^95^. We retained the remaining unplaced sequences for downstream analysis.

To validate the NRSs extracted by the strategy in this study, we extracted NRSs from *de novo* assembly of HG002 with 15-fold ONT reads and compared them with the NRSs extracted from the HG002 assembly used as GIAB benchmark dataset^14^. In addition, we *de novo* assembled the genomes of 10 samples in this study sequenced by PacBio HiFi reads and then extracted NRSs (**Table S3**). We then detected how many NRSs from these 10 samples with ONT reads can be validated by NRSs from assemblies of PacBio HiFi data.

To analysis the hotspot of NRSs located in the genome, we applied the function “hotspotter” from the primatR package (https://github.com/daewoooo/primatR) with parameters “bw=200000, num.trial=1000”. A p value was calculated by comparing the increased density of NRS locations around the genome with the density of a randomly subsampled set of the genome.

### Non-redundant NRS of the population

We applied Jasmine^96^ v1.1.0 to generate non-redundant sequences for placed and unplaced NRSs. For the placed NRSs, we first calculated the median anchored position in the reference genome for the upstream and downstream of this sequence. Then we used Jasmine to combine the placed NRSs using parameters “--output_genotypes -- ignore_strand --keep_var_ids”. For all pairs of sequences in each cluster, the distance of each paired sequences also restricted to 250 bp. For each cluster, we used Kalign^97^ v3.3 to perform multiple sequence alignment and calculated the score of each sequence (match +2, mismatch −1, gap opening −0.5). Then we selected the sequence with highest score as the representative sequence. For all unplaced NRSs, we performed all-versus-all alignments using minimap2 with parameters “-DP -t threads unplaced.fa unplaced.fa > aligned.paf”. The alignment pair with the alignment length ≥ 200 bp and sequence identity ≥ 90% were used for downstream analysis, while the sequence without intersections to other sequences was considered as non-redundant sequence. For the alignment pair, we removed the shorter one if the alignment length covered at least 80% of one sequence. And the unplaced sequences existed in only one individual were removed. Consequently, we obtained the non-redundant NRSs for the whole population. We divided the NRSs into four categories based on the AF of NRS in the whole population: singleton (allele count = 1), polymorphic (allele count ≥ 2 and AF < 0.5), major (AF ≥ 0.5 and AF < 1) and shared (AF = 1).

### Graph-based pangenome of NRS and genotyping

To genotype the NRSs, we first built a graph-based pangenome of NRSs. In short, the placed NRSs and the reference genome GRCh38 were used to build a graph reference using vg with parameters “vg construct -a -f -p -S -m 32”. And the unplaced NRSs were added to the end of pangenome. The placed NRSs were genotyped for all individuals using vg toolkit^98^ v1.33.1 and GraphAligner^38^ v1.0.13, which was pan-genome graph-based SV genotyper and tool for aligning long reads to genome graphs, respectively. First, long reads of each individual were aligned to the graph reference with GraphAligner using parameter “-x vg”. Then, vg was used to genotype the NRSs according to the long-read alignment. If the NRSs were extracted from an individual, the genotype of NRSs for this individual should be “0/1” or “1/1”. We evaluated recall rate of genotyping based on presence and absence information of NRSs. For the short-read sequences from Illumina platform, we used vg giraffe aligned the reads to the graph genome with parameters “-p -b default --rescue-algorithm dozeu”.

### Comparison of NRS to other human genomes and pan-genomes

Human assembly genomes from different ancestries were reported, such as T2T-CHM13^2^, HX1^10^, AK1^99^, KOREF^100^, HuRef^101^, NA12878^9^ and NA19240^102^ (**Fig. 2a**). In addition, some human pan-genomes, such as Chinese HUman Pan-genome Analysis (HUPAN)^6^, African pan-genome (APG)^103^, Icelander non-repetitive non-reference sequences (Icelander-NRNR)^5^, 1,000 Swedish genomes (PanSwe)^104^, Swedish genomes (TwoSwe)^11^, Mix-17NUIs^105^ and HGD-NUIs^106^, had been reported by large-scale whole-genome sequencing using NGS data^3^ (**Fig. 2b**). Reciprocal alignments between our NRSs and these assemblies were conducted using minimap2 with parameters “-x asm20 -t threads nonref.fa genome.fa > aligned.paf” and retained alignments with an overall identity ≥ 90% and coverage ≥ 80% of the NRSs^103^. The alignment was regarded as reliable if aligned length ≥ 200 bp with the aligned identity ≥ 90% and aligned coverage ≥ 80% of this sequence.

### Annotation of non-redundant NRS

To reduce the false rate of the protein-coding gene annotation, we first masked the repeat sequences as “N” using method mentioned above. Then we downloaded all expressed sequence tags (ESTs) of human from Ensembl and human protein sequences from NCBI (v2020-3-17). The redundant sequences in ESTs and protein sequences were independently removed using CD-HIT^107^ v4.8.1 with parameters “cd-hit-est -i est.fa -o est.cdhit.fa -c 0.9 -n 8 -d 0 -M 0 -T threads” and “cd-hit -i protein.fa -o protein.cdhit.fa -c 0.9 -n 5 -d 0 -M 0 -T threads”, respectively. Then, protein-coding genes for non-redundant NRSs were predicted from the repeat-masked non-reference genome using MAKER2^108^ v2.31.1. In this process, SNAP^109^ (v2006-07-28) was trained for two rounds based on EST sequences, and *ab initio* gene prediction was performed using Augustus^110^ v3.3.3 based on human model. Furthermore, we retained the predicted transcript with length > 150 bp. We also used models with an annotation edit distance (AED) of ≤ 0.5 to help remove junky models^111^. To get reliable predicted genes, we used blastn and blastx to search all open reading frames against all sequences in NCBI nt and nr databases, respectively, to determine whether the gene was conserved in other primates or elsewhere in the mammalian genome. The thresholds of *e*-values for blastn and blastx were set as 1 ×10^−15^ and 1 ×10^−7^, respectively. We then annotated the domains that matched either the Pfam protein families database^26^ or the NCBI Conserved Domain Database (CDD)^25^. The predicted gene without intron was regarded as pseudogene.

### NRS associated expression quantitative trait loci (eQTLs)

To test the impact of NRSs to gene expression levels, we first quantified the association between gene expression levels and NRSs within a 1 Mb window, centred around the gene’s annotated transcription start site^17^, by employing a panel of 451 individuals from GEUVADIS Consortium^42^. Based on the graph pangenome of NRSs constructed in this study, we genotyped the NRSs for the panel of 451 individuals from the whole-genome NGS data. And 7,244 NRSs with genotype information at MAF > 0.05 were selected. Subsequently, we quantified the transcript-level expression of this panel based on RNA data using graph-based method via vg mpmap and rpvg (https://github.com/jonassibbesen/rpvg) according to the previously reported pipeline^112^. To identify NRS associated eQTLs from the panel of 451 individuals, we conducted PCA analysis from the genotype matrix of NRSs in this panel, followed by the association analysis between transcript-level expression and genotypes of NRSs within a 1 Mb window, implemented by fastQTL^113^ v2.165. We next used the Benjamini-Hochberg procedure to identify all NRS eQTL pairs at 5% FDR.

### Population stratification and local adaption

To assess the population stratification of Africans (AFRs), Americans (AMRs) and East Asians (EASs), we performed principal component analysis (PCA) using EIGENSOFT^114^ v7.2.1 using 4,685 NRSs (MAF > 0.05) fitting HWE. Then we calculated population branch statistics (PBS) for subpopulations using PBScan^115^ v2020-03-16. All the NRSs that were polymorphic within subpopulations were used for further analysis. And we regarded the rank of 99.9% as the threshold for evidence of departure from neutrality. We calculated PBS using AFR, AMR and EAS, and 13,518 NRSs (MAF > 0.01) were used. The PBS thresholds for AFR and EAS, and AMR and EAS were 0.62 and 0.42, respectively. SVs with PBS score above the threshold within continuous 1 Mb were combined as an independent signal. To obtain high-confidence loci, we filtered out the loci that may have batch effects which may result from the datasets across multiple platforms. We performed a chi-squared test for the NRS genotypes within the population, such as the genotypes from PacBio CLR and HiFi in AFRs, and the ones from ONT and PacBio HiFi in EASs. And p values were corrected by the Benjamini-Hochberg method. And the loci with corrected q value < 0.05 were defined as batch effect and were excluded from the PBS result.

### Genotype and phenotype association analysis

In this study, 5,643 genotyped NRSs with minor allele frequency (MAF) > 0.05 in 327 individuals with clinical phenotypes were used for the analysis. The genome-wide association study (GWAS) was performed using PLINK^116^ v1.90b4 with linear regression under an additive genetic model for the quantitative traits, and age, sex, body mass index (BMI), and the first two principal components were included as covariates. When applying BMI GWAS, BMI itself was excluded from covariates. The association test for case-control was conducted using logistic regression module. And the significant threshold was set to be 8.9×10^−6^ through Bonferroni correction (0.05/5,643)^117^.

### SNP detection and linkage disequilibrium analysis

For the 405 samples in our previous study^18^, we detected the SNPs across the genome for each sample using longshot^118^ based on the BAM files against the reference GRCh38. To obtain the high-quality SNPs, we filtered the result with supported reads ≥ 8 and Qual ≥ 20. Meanwhile, 105,440 SNPs associated with the phenotypes in GWAScatalog^74^ r2020-03-08 were extracted as the target SNP dataset. Subsequently, we constructed the matrix of the target SNP and 405 samples. If the SNP was not called in the target locus for the sample, the genotype was recoded as the allele of homologous reference (0/0). The linkage disequilibrium between the target SNP dataset and NRSs detected in this study was calculated using PLINK.

### Statistical analysis

For each NRS, we conducted Fisher’s exact test to calculate the Hardy-Weinberg *P* values. NRSs with p value < 0.0001 were considered as HWE-failed. Wilcoxon signed-rank test was conducted for the read mapping rates derived from linear reference genome and pamgenome. We performed a chi-squared test for the genotypes from different platforms within the population. And *P* values were corrected by the Benjamini-Hochberg method. The GO annotation files, including GO_Molecular_Function_2018, were downloaded from Enrichr^119^ website (https://maayanlab.cloud/Enrichr/). And the GO enrichment analysis was performed using Fisher’s exact test, followed by *P* value correction using Benjamini-Hochberg method. The statistical tests used were described throughout the article and in the figures. In the boxplots, the upper and lower hinges represented the first and third quartile. The whiskers extended to the most extreme value within 1.5 times the interquartile range on either end of the distribution, and the center line represented the median.

## Data availability

Sequencing data for all the 539 individuals in this study are publicly available. And the detail information of these datasets was list in **Table S2**. The sequences and genotypes of the NRSs are publicly available at the National Genomics Data Center (NGDC), China National Center for Bioinformation (CNCB) with project accession number PRJCA007976 (https://ngdc.cncb.ac.cn/bioproject/browse/PRJCA007976). The sequences and genotypes of the placed NRSs are available with accession number GVM000324 (https://ngdc.cncb.ac.cn/gvm/getProjectDetail?project=GVM000324). And the sequences of the unplaced NRSs are under the accession number GWHBHSK00000000 (https://ngdc.cncb.ac.cn/gwh/Assembly/24529/show).

## Code availability

The codes of pipeline GraphNRS in this study are publicly available via GitHub repository (https://github.com/xie-lab/GNRS).

## Acknowledgements

We would like to thank all the data contributors who make this research possible, particularly the Genome in a Bottle Consortium, Telomere-to-Telomere Consortium, and the Human Pangenome Reference Consortium. We also thank the Center for Precision Medicine at Sun Yat-sen University for the long-term support.

## Funding

This project was supported by National Key R&D Program of China (2019YFA0904400, Z.X.), National Natural Science Foundation of China (31829002, Z.X.).

## Author information

These authors contributed equally: Zhikun Wu and Tong Li.

## Authors and Affiliations

State Key Laboratory of Ophthalmology, Zhongshan Ophthalmic Center, Sun Yat-sen University, Guangzhou, China

Zhikun Wu, Tong Li, Zehang Jiang, Jingjing Zheng, Yizhi Liu & Zhi Xie

MOE Key Laboratory of Metabolism and Molecular Medicine, Department of Biochemistry and Molecular Biology, School of Basic Medical Sciences and Shanghai Xuhui Central Hospital, Fudan University, Shanghai, China

Yun Liu

## Contributions

Z.X conceived and supervised the study. ZK.W and Z.X designed the study. T.L and ZK.W analysed the data. All the authors interpreted the data. ZK.W, Z.X, T.L and ZH.J wrote the manuscript. All authors read and approved the manuscript.

## Competing interests

The authors declare that they have no competing interests.

## Notes

### Competing Interest Statement

The authors have declared no competing interest.

