## Supplemental Figures for "Graph pangenome reveals functional, evolutionary, and phenotypic significance of human nonreference sequences"

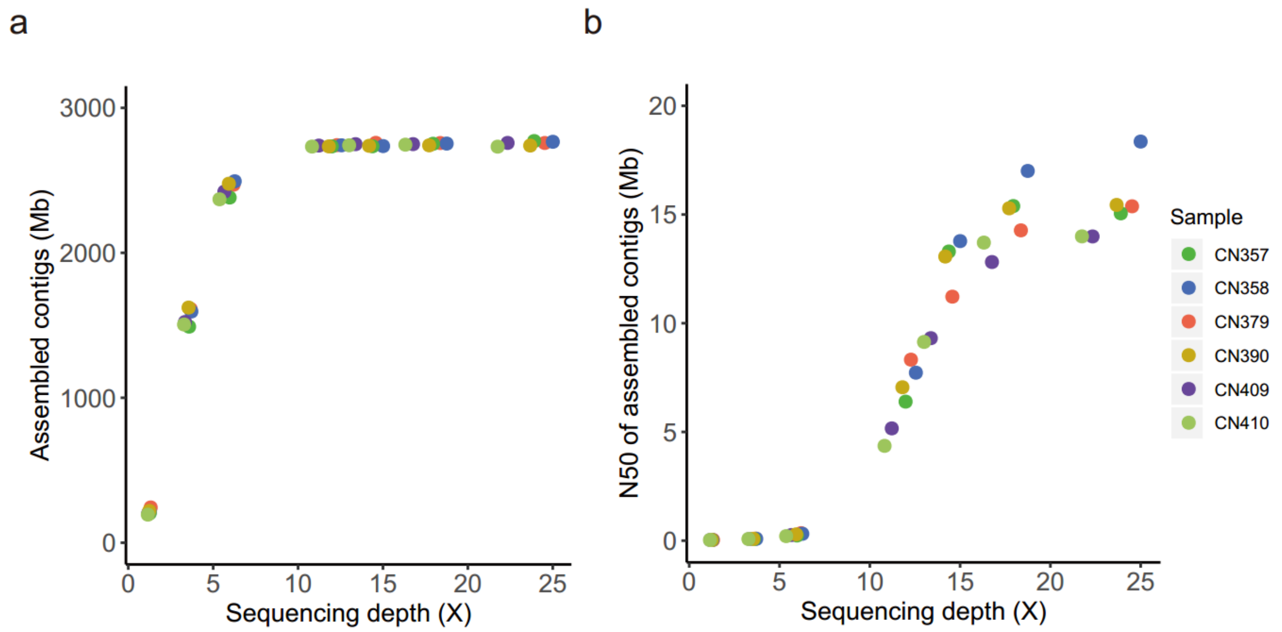

**Figure S1. Lengths of assemblies versus different sequencing depths**

**a**, Total lengths of assemblies versus different sequencing depths.

**b**, N50 length of assembled contigs versus different sequencing depths.

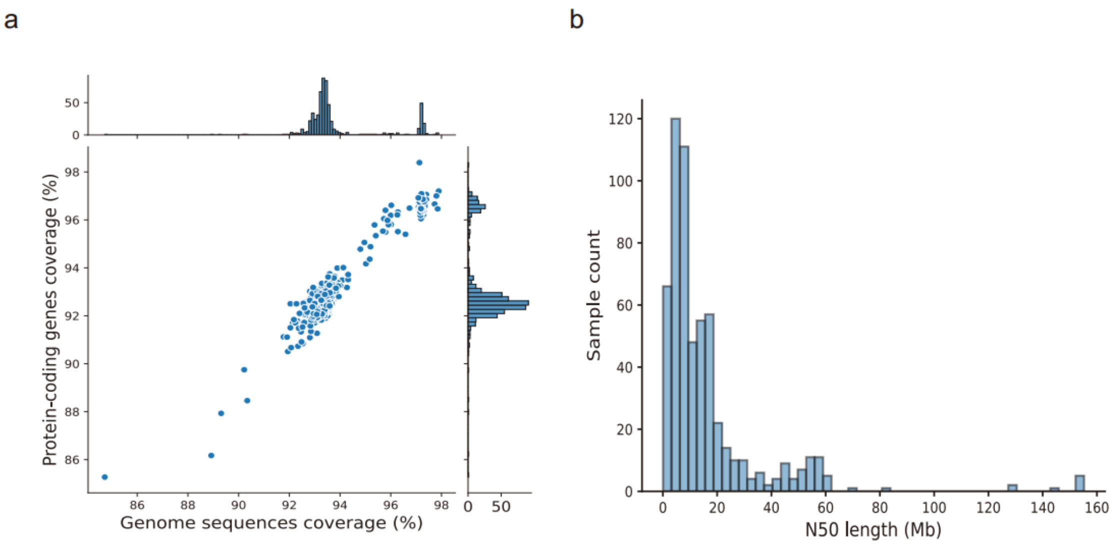

**Figure S2. Evaluation of *de novo* assembled genomes**

**a**, Distribution of genome sequence coverage and protein-coding gene coverage.

**b**, Distribution of N50 length of assembled contigs.

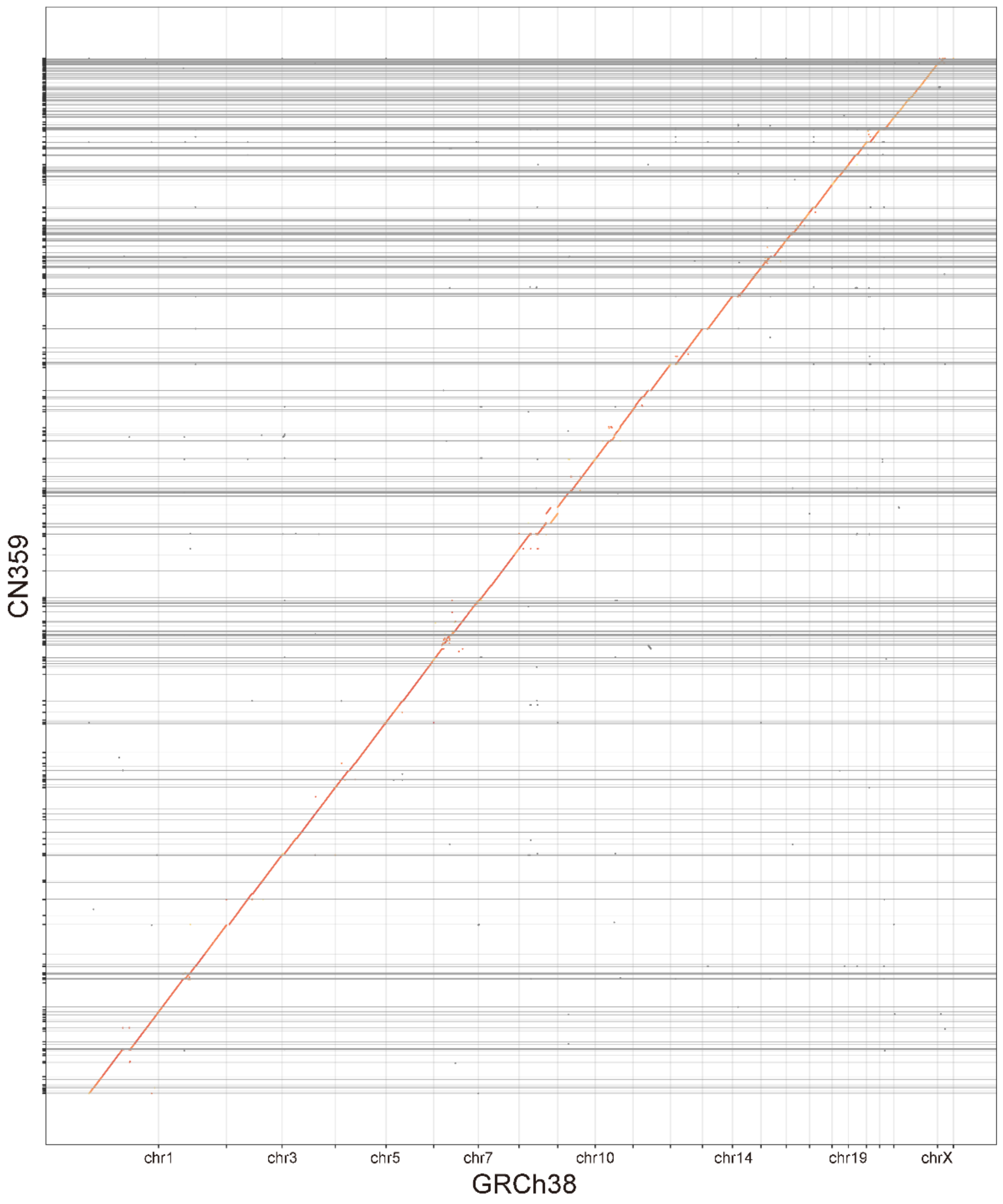

**Figure S3. Macrosynteny between assembled contigs and the reference genome CRCh38**

The x and y axes are the coordinates of the reference genome GRCh38 and the assembly in this study, respectively.

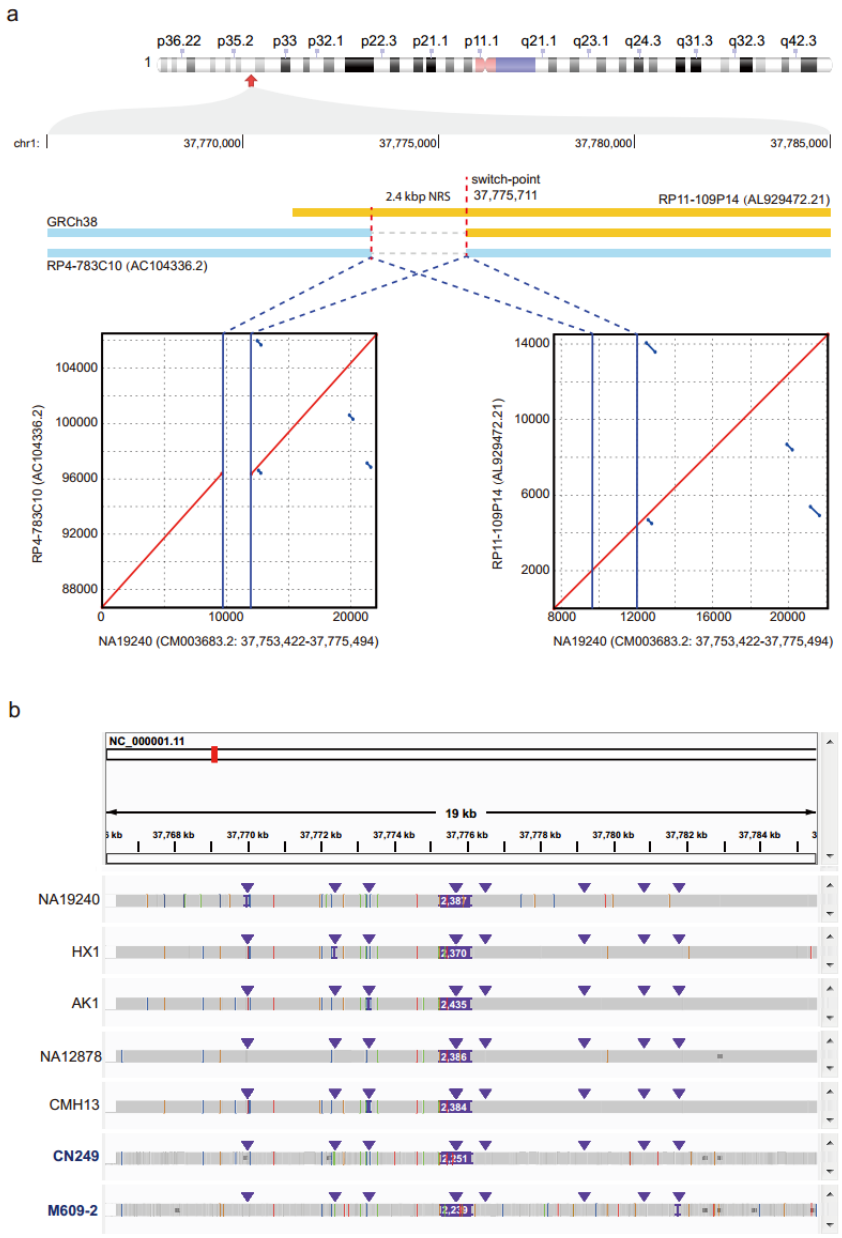

**Figure S4. An example of mis-assembly from GRCh38**

**a**, A 2.4 kb NRS was precisely anchored at the switch-point of BACs RP4-783C10 (AC104336.2) and RP11-109P14 (AL929472.21). Compared to NA12940 (x-axis), the deleted sequence of RP4-783C10 (AC104336.2) (y-axis) resulted in a missing sequence in final assembly of GRCh38 in the switch-point of these two BACs.

**b**, The diagram shows that several published genomes contain a 2.4 kb NRS at the switch-point of GRCh38. The sample name in blue indicates genome assembly generated in this study.

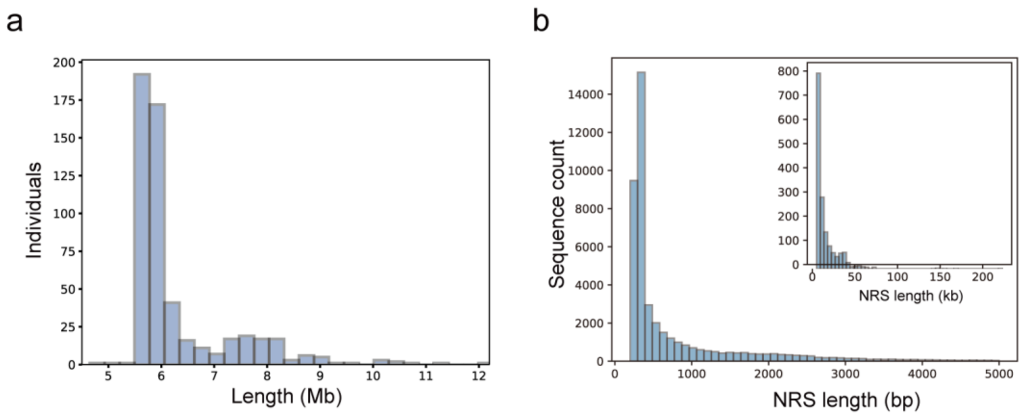

**Figure S5. Length distribution of the NRSs.**

**a**, Length distribution of the extracted NRSs for the 539 *de novo* assembled genomes.

**b**, Length distribution of the non-redundant NRSs for the whole population.

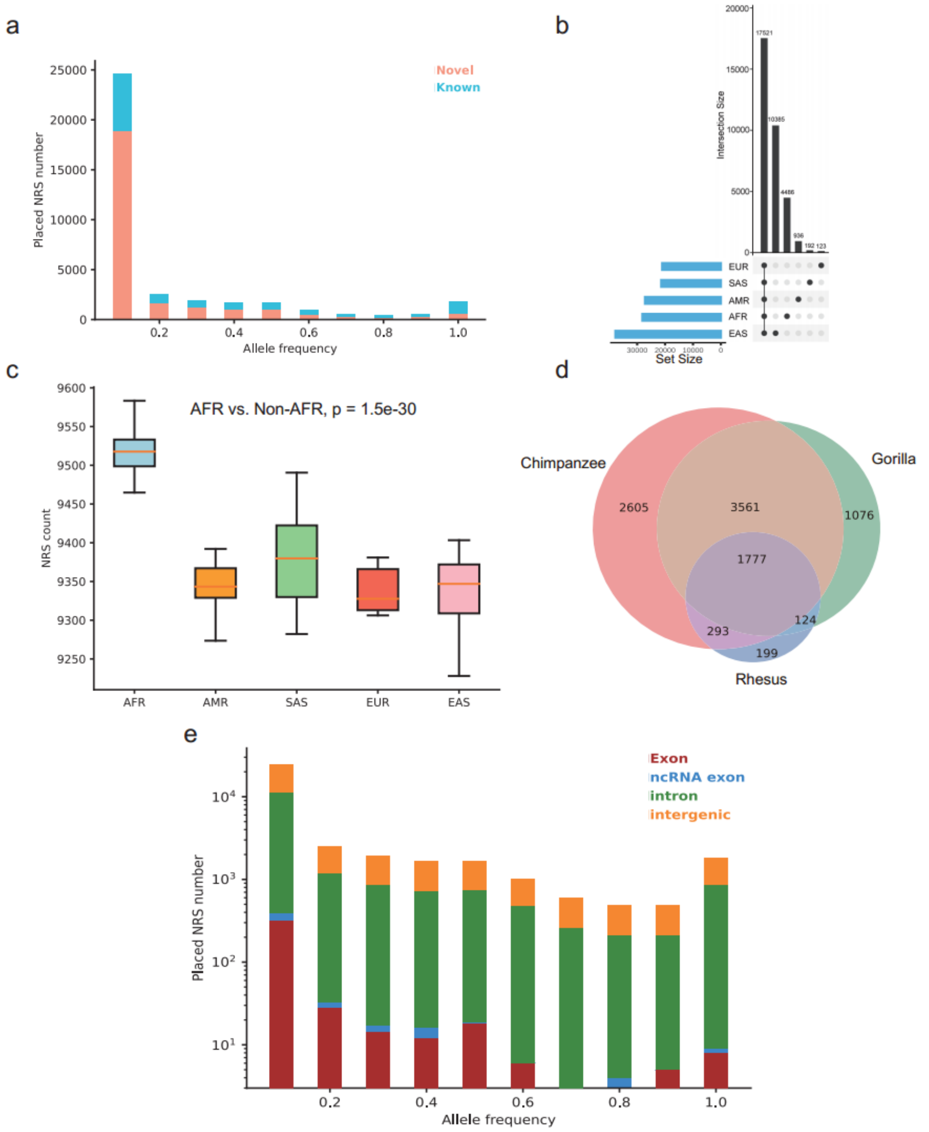

**Figure S6. Characterization of NRSs for the whole population**

**a**, Allele frequency of the non-redundant NRSs.

b, Summary of common and specific NRSs in diverse populations.

**c**, Boxplots show the NRS counts for different populations. The NRS counts across different platforms were normalized using z-score. AFR: African, AMR: American, SAS, South Asian, EUR, European, EAS, East Asian. The center line in the box indicates the median, the lower and upper hinges indicate the first and third interquartile range (IQR). The lower and upper whiskers show the values greater than 25th quartile minus 1.5 × IQR and less than 75th quartile plus 1.5 × IQR, respectively. Where data beyond these ranges are shown as individual points.

**d**, The number of overlapped NRSs among three non-human primate genomes.

**e**, The gene feature annotation of placed NRSs with different allele frequencies.

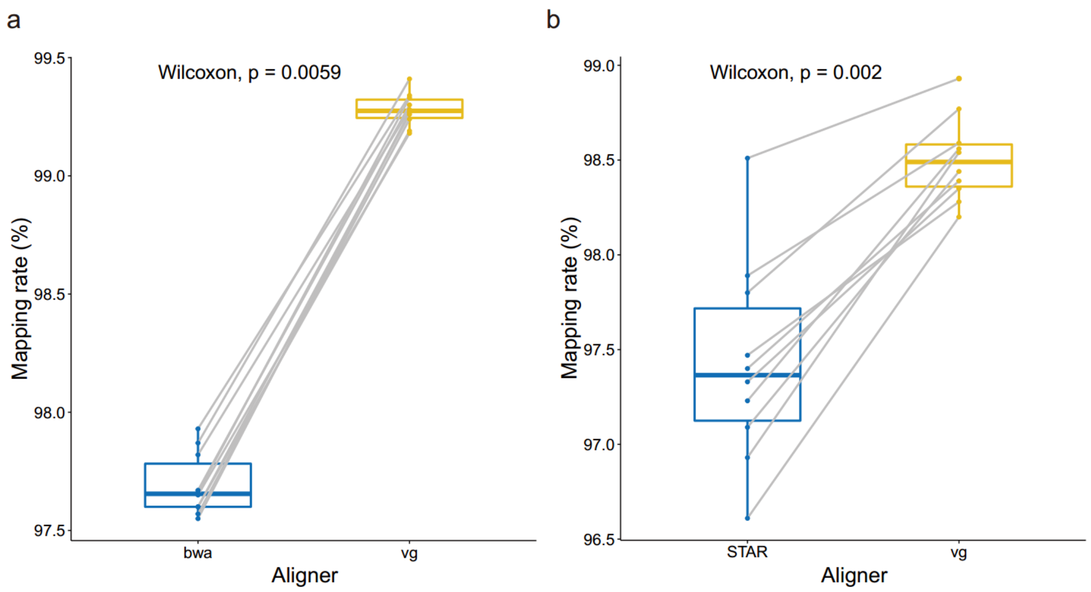

**Figure S7. Mapping rate improvement of short-read sequencing data**

a, The mapping rate improvement of DNA from short-read sequencing platform, bwa and vg are aligners against the conventional linear genome GRCh38 and graph pangenome in this study, respectively.

b, The mapping rate improvement of RNA from short-read sequencing platform, STAR and vg are aligners against the conventional linear genome GRCh38 and graph pangenome, respectively.

Wilcoxon signed-rank test was conducted for the mapping rates of ten samples.

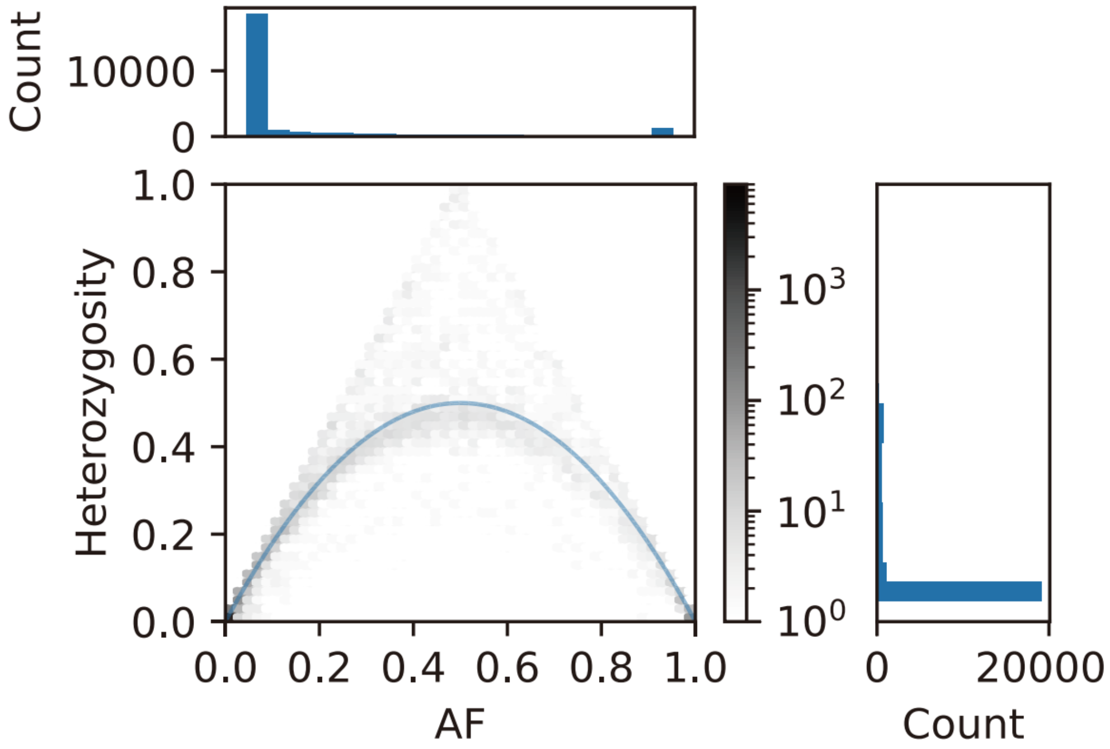

**Figure S8. Genotyping information of NRSs.**

The plot shows the relationship between allele frequency (AF) and heterozygosity of the genotypes of NRSs derived from the graph pangenome.

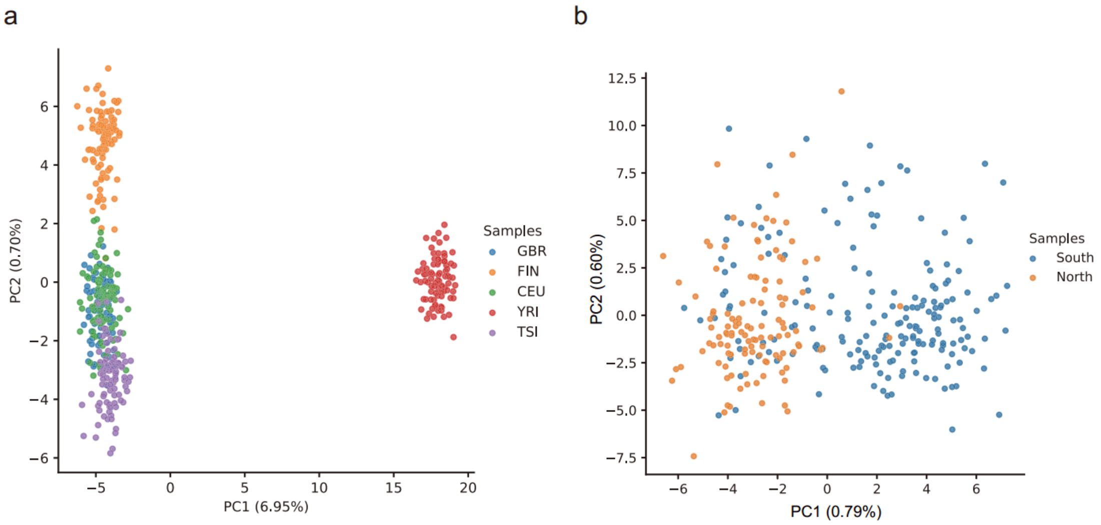

**Figure S9. Population stratification analysis based on genotypes of NRSs**

a, The principal component analysis (PCA) based on NRSs from short-read sequencing data detects population stratification for Genetic European Variation in Disease (GEUVADIS) consortium, which consist of four European-ancestry and one African-ancestry populations. GBR: British, FIN: Finnish, CEU: Utah residents (CEPH), TSI: Toscani, YRI: Yoruba. The values in parentheses indicate the genetic variations explained by the first two PCs.

b, PCA based on NRSs from long-read sequencing data between southern and northern Chinese populations in this study.

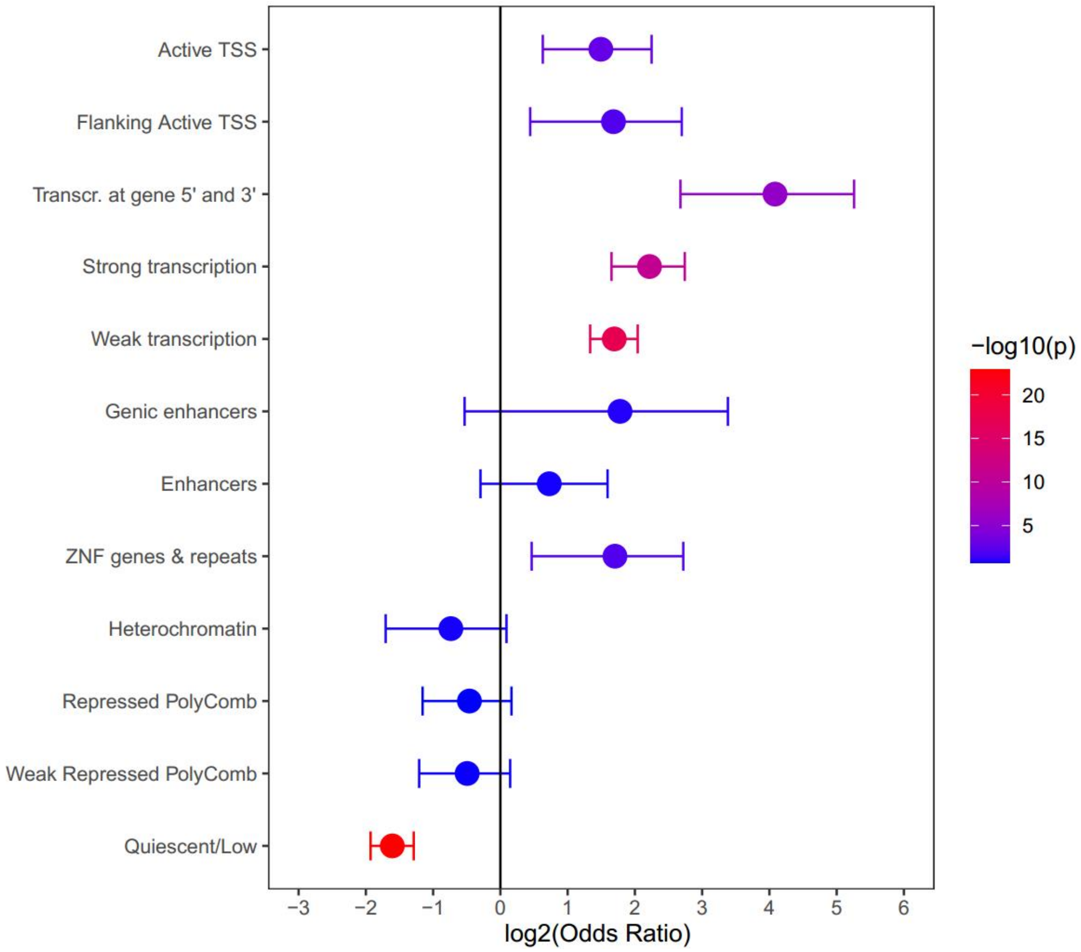

**Figure S10. Enrichment or depletion of eQTL-associated NRSs**

The enrichment or depletion of eQTL-associated NRSs that intersected with the epigenetic states by the Roadmap Epigenetics Consortium (REC). The Fisher's exact test was conducted, and p values were corrected using Benjamini-Hochberg method.

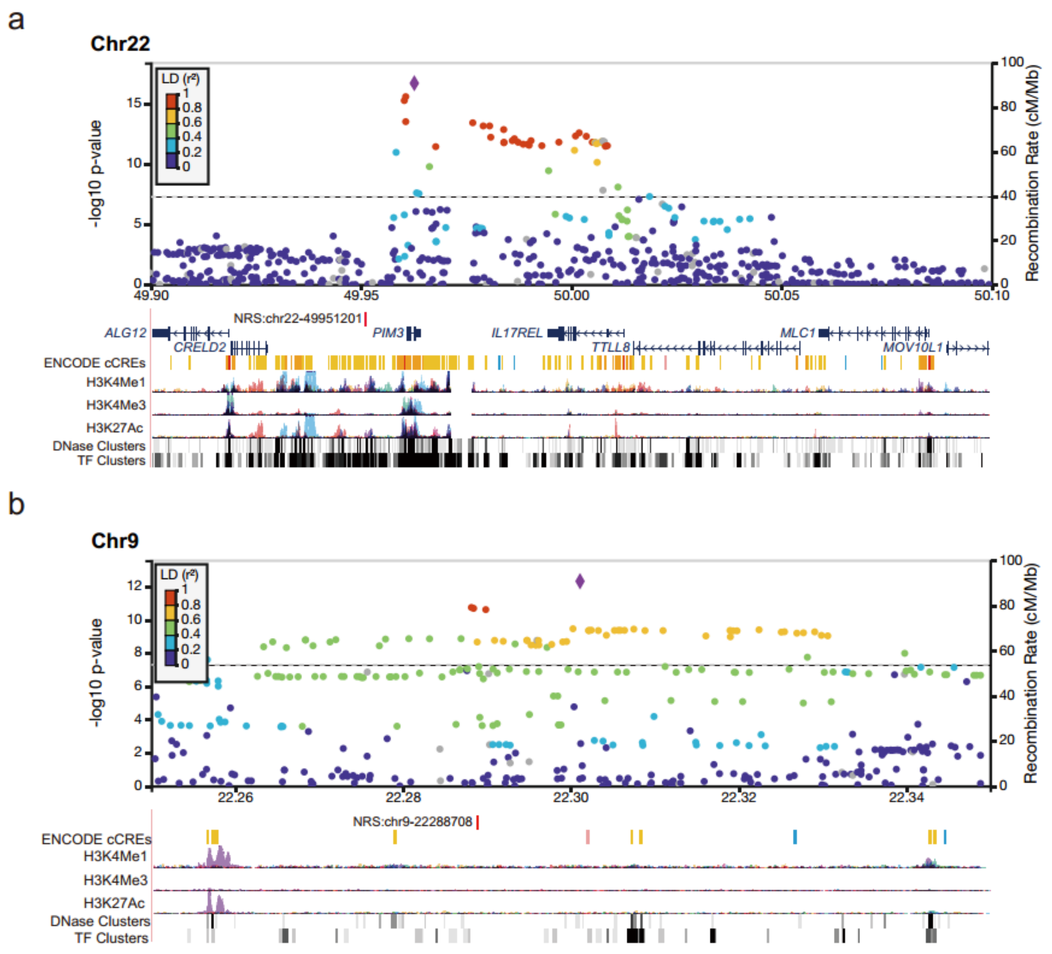

**Figure S11. The regional manhattan plot for SNPs associated with diabetes.**

a, The top signal was significantly associated with diabetes ($P$ = 1.8 $\times 10^{-17}$). The NRS is 9.6 kb upstream of *PIM3* and intersected with H3K27Ac, H3K4Me1 and TF clusters.

b, The top signal was significantly associated with diabetes ($P$ = 4.7 $\times 10^{-13}$). There are three SNPs in high LD ($r^2$ > 0.8) with top signal of diabetes. And the NRS (red vertical line) located in the region of three SNPs.

The pairwise $r^2$ values were from the 1000 Genome Phase3 (EAS) reference panel.
